# *Salmonella enterica* serovar Typhi uses two type 3 secretion systems to replicate in human macrophages and to colonize humanized mice

**DOI:** 10.1101/2023.06.06.543980

**Authors:** Meagan Hamblin, Ruth Schade, Ramya Narasimhan, Denise M. Monack

**Affiliations:** Department of Micrology and Immunology, Stanford University School of Medicine, Stanford, Califonia, USA

## Abstract

*Salmonella enterica* serovar Typhi (*S.* Typhi) is a human-restricted pathogen that replicates in macrophages. In this study, we investigated the roles of the *S.* Typhi Type 3 secretion systems (T3SSs) encoded on *Salmonella* Pathogenicity Islands (SPI) -1 (T3SS-1) and -2 (T3SS-2) during human macrophage infection. We found that mutants of *S*. Typhi deficient for both T3SSs were defective for intramacrophage replication as measured by flow cytometry, viable bacterial counts, and live time-lapse microscopy. T3SS-secreted proteins PipB2 and SifA contributed to *S.* Typhi replication and were translocated into the cytosol of human macrophages through both T3SS-1 and -2, demonstrating functional redundancy for these secretion systems. Importantly, an *S*. Typhi mutant strain that is deficient for both T3SS-1 and -2 was severely attenuated in the ability to colonize systemic tissues in a humanized mouse model of typhoid fever. Overall, this study establishes a critical role for *S.* Typhi T3SSs during its replication within human macrophages and during systemic infection of humanized mice.

**Importance:** *Salmonella enterica* serovar Typhi is a human-restricted pathogen that causes typhoid fever. Understanding the key virulence mechanisms that facilitate *S.* Typhi replication in human phagocytes will enable rational vaccine and antibiotic development to limit spread of this pathogen. While *S.* Typhimurium replication in murine models has been studied extensively, there is limited information available about *S.* Typhi replication in human macrophages, some of which directly conflicts with findings from *S.* Typhimurium murine models. This study establishes that both of *S.* Typhi’s two Type 3 Secretion Systems (T3SS-1 and -2) contribute to intramacrophage replication and virulence.

## Introduction

Typhoid fever is a transmissible disease caused by the bacterium *Salmonella enterica* serovar Typhi (*S*. Typhi). Before widespread antibiotic use, over 50% of typhoid cases resulted in serious complications, with a mortality rate of over 10% (1). Outbreaks of multi- and extensively-drug resistant typhoid are increasing in magnitude and frequency, and climate change will likely exacerbate transmission, underscoring the need to identify new antimicrobial targets (2, 3).

The molecular mechanisms underlying typhoid pathology are poorly understood due to the lack of experimental animal models, as *S*. Typhi is human-restricted. By studying the related pathogen *Salmonella enterica* serovar Typhimurium (*S*. Typhimurium), which causes a typhoid-like disease in mice, researchers have identified many molecular mechanisms underlying *S.* Typhimurium virulence. However, these two serovars cause distinct disease states during human infection. Although both pathogens can replicate inside phagocytes, *S*. Typhimurium infection is usually restricted to the human gastrointestinal tract (4). On the other hand, *S*. Typhi infection frequently disseminates systemically in humans, and can result in internal bleeding, hepatic dysfunction, splenomegaly, intestinal perforation, and typhoid encephalopathy (3, 4). Furthermore, the ability of *Salmonella* to proliferate within phagocytic cells, including macrophages, is a hallmark of systemic disease (5).

*S*. Typhi and *S*. Typhimurium have differentially evolved to encode distinct virulence factors (6–8). Researchers have identified many genes essential for *S*. Typhimurium pathogenesis in mice, some of which are absent, functionally null, or dispensable in *S*. Typhi pathogenesis studies. One example is the type 3 secretion system encoded in *Salmonella* pathogenicity island 2 (T3SS-2) which is important for establishing and maintaining a membrane compartment, the *Salmonella*-containing vacuole (SCV), that *Salmonella* replicates in (9). Inhibition or deletion of T3SS-2 renders *S*. Typhimurium unable to replicate in mouse macrophages (10, 11). In contrast, *S.* Typhi replication is not dependent on T3SS-2 in human macrophages (12, 13). However, *S.* Typhi has been shown to express T3SS-2 during human macrophage infection (14). Additionally, the T3SS-2 encoded by *S.* Typhi translocates bacterial effector proteins into host cells during intracellular infection (15).

*S*. Typhi has an additional T3SS encoded in SPI-1 (T3SS-1). T3SS-1 contributes to invasion and cytosolic replication in epithelial cells (15). Furthermore, T3SS-1 was recently shown to be essential for *S*. Typhi survival in human stem-cell derived macrophages (16). Based on these previous findings, we hypothesized that both T3SSs in *S.* Typhi contribute to replication in human macrophages. To characterize the relative contributions of *S.* Typhi T3SSs to virulence, we created strains deficient for either T3SS-1 or -2 and a strain lacking both T3SSs and characterized the kinetics of replication within human-derived macrophages and in humanized mice.

## Results

### *Salmonella* Typhi uses both T3SS-1 and T3SS-2 to replicate in human macrophages

To assess the relative contributions of T3SS-1 and -2 to *S.* Typhi replication within macrophages, we infected PMA-differentiated THP-1 macrophages, a human-derived promonocytic cell line commonly used to model *Salmonella* infection (12, 17–19). To measure *S.* Typhi replication in THP-1 macrophages, we used a previously published method of quantifying fluorescence dilution (pFCcGi), a dual-fluorescence tool which permits direct assessment of intramacrophage bacterial replication (11, 20). By flow cytometry, the wild-type (WT) *S*. Typhi Ty2 strain replicated 8- to 11-fold over 24 h (Fig. 1A). In contrast, an isogenic *S*. Typhi Δ*phoP* strain, which is deficient for a virulence transcriptional regulatory protein and is unable to survive within macrophages, did not replicate in THP-1 macrophages (Fig. 1A), which is consistent with previous studies (12, 13). To test the roles of the T3SS-1 and T3SS-2 in *S*. Typhi replication within human macrophages, we deleted InvA (Δ*invA*) or SsaV (Δ*ssaV*), the export gates in T3SS-1 and T3SS-2, respectively. The *S*. Typhi Δ*invA* strain replicated in THP-1 macrophages to the same extent as WT *S*. Typhi (8-fold) whereas the *S.* Typhi Δ*ssaV* strain replicated slightly less than the WT strain (7-fold) (Fig. 1A). To test the possibility that both T3SSs contribute to intramacrophage replication, a double knock-out *S.* Typhi Δ*invA*Δ*ssaV* (T3SS-null) strain was constructed and intramacrophage replication was measured. In contrast to the single T3SS-deficient strains, the strain that is deficient for both T3SSs had a severe defect in THP-1 macrophages (3 to 5-fold replication), nearly phenocopying the Δ*phoP* strain (Fig. 1A). To validate the fluorescence dilution results, viable bacteria were enumerated at 2 and 24 hours post-inoculation (h.p.i.). by CFU plating (Fig. S1A). In agreement with the fluorescence dilution assay results, WT *S.* Typhi replicated to significantly higher levels (∼10-fold) compared to the Δ*phoP* strain (Fig. S1B). In contrast, the strains that are deficient for one of the T3SSs (the Δ*invA* and Δ*ssaV* strains) replicated to the same extent as the WT *S.* Typhi strain. Strikingly, the *S.* Typhi strain lacking both T3SSs (T3SS-null strain) had a significant replication defect compared to the WT *S.* Typhi strain (Fig. S1B). Importantly, the WT and T3SS-deficient bacterial strains were not defective for uptake by THP-1 macrophages (Fig. S1A), or for growth in a defined minimal medium (Fig. S1C), further validating an intramacrophage-specific replication defect. Finally, the levels of cell death induced during 24-hour infections with WT or T3SS-deficient bacterial strains were low and similar for all strains (Fig. S1D), ruling out host cell death as a potential confounding factor.

**Figure 1.**
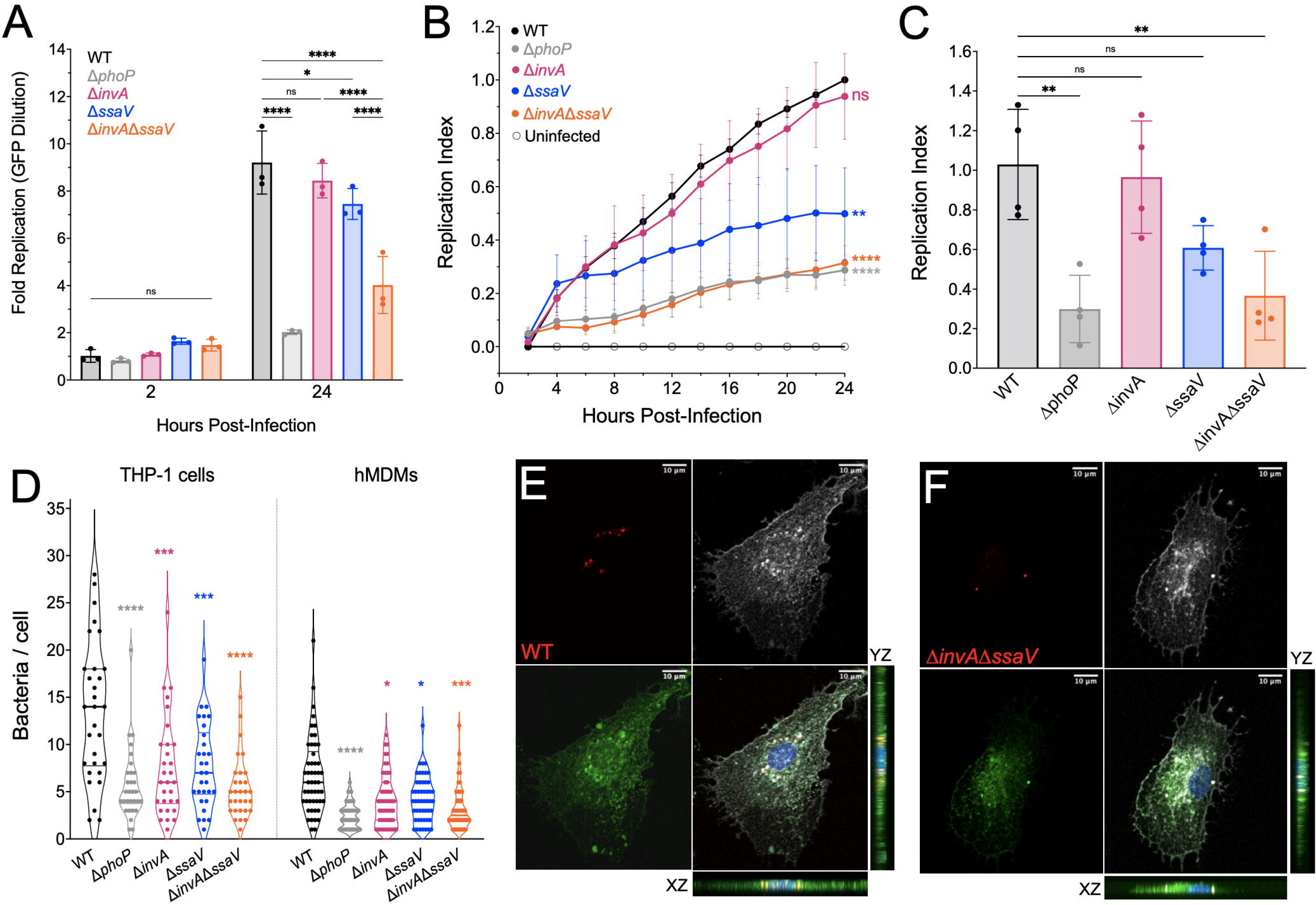
*S.* Typhi uses both T3SS-1 and T3SS-2 to replicate in human macrophages. **A.** Replication of *S.* Typhi in THP-1 macrophages by fluorescence dilution (pFCcGi) at 2 and 24 h.p.i.. Statistical significance by ANOVA. One biological replicate representative of 3 replicates Dots: technical replicates. Bars: mean. Error: SD. **B.** Replication of *S.* Typhi in hMDMs measured by fluorescence dilution (pFCcGi) by time-lapse microscopy during 24 h infection. Statistical significance compared to WT at 24 h.p.i. by ANOVA. Dots: mean of 6 biological replicates. Error: SEM. **C.** Replication of *S.* Typhi in hMDMs measured by fluorescence dilution (pFCcGi) at 20 h.p.i. by time-lapse microscopy. Statistical significance compared to WT by ANOVA. Bars: mean of 4 biological replicates, 3 technical replicates each. Error: SD. **D.** Number of bacteria per cell in fixed macrophages at 16 h.p.i. Statistical significance by ANOVA. Dots: number of bacteria counted in one cell. Violin outline: population distribution. Thick line: median. Thin line: upper and lower quartiles. **E.** WT-infected hMDMs, 63X. Maximum intensity projection of Z-stack images taken with confocal microscope. Blue= nuclei, Green = LAMP-1, White = actin, Red = *Salmonella*. Scale: top right. **F.** Δ*invA*Δ*ssaV*-infected hMDMs, 63X. Maximum intensity projection of Z-stack images taken with confocal microscope. Blue= nuclei, Green = LAMP-1, White = actin, Red = *Salmonella.* Scale: top right. For all; ns = p-value > 0.05, *≤0.05, ** ≤0.01, *** ≤0.001, **** ≤0.0001

To compare the kinetics of *S.* Typhi intramacrophage replication, we then performed live-cell imaging and quantified fluorescence dilution (pFCcGi) over 24 hours for each strain relative to WT (Videos S1-2) (11, 20). For each experiment, we obtained the ratio of mCherry to GFP signal within macrophages (Fig. S1E), subtracted background signal measured in uninfected wells, then normalized to that of the wild-type strain at 24 h.p.i. to obtain a “Replication Index”. Throughout the infection, WT and Δ*invA S*. Typhi strains replicated to similar levels in THP-1 macrophages (Fig. 1B). Although the *S*. Typhi Δ*ssaV* strain replicated less than the WT strain at later timepoints, the T3SS-null strain had a significant replication defect throughout the time course, nearly phenocopying the Δ*phoP* control (Fig. 1B). To confirm that T3SS-dependent replication also occurs in primary human macrophages, we infected human blood monocyte-derived macrophages (hMDMs) and performed time-lapse microscopy to quantify replication at several timepoints during infection (Videos S3-4, Fig. S1F). Throughout infection, the T3SS-null mutant had a severe replication defect compared to the WT strain, phenocopying the Δ*phoP* strain, whereas the single Δ*invA* and Δ*ssaV* knockout *S*. Typhi strains had intermediate replication defects (Fig. 1C). Finally, we counted the number of intracellular bacteria per THP-1 macrophage or hMDM by confocal microscopy at 16 h.p.i. Although THP-1 macrophages contained a higher overall abundance of *S*. Typhi compared to hMDMs, the relative contributions of T3SS-1 and -2 to the abundance of bacteria inside hMDMs was similar to what was observed in THP-1 macrophages (Fig. 1D).

To confirm that gentamicin-protected *S.* Typhi were intracellular, hMDMs infected with either WT or T3SS-null bacteria were stained with an antibody to the endosomal membrane marker LAMP-1 and phalloidin to stain polymerized actin and analyzed by confocal microscopy. Both WT and T3SS-null *S.* Typhi colocalized with LAMP-1 and actin, suggesting that the gentamicin-protected bacteria reside within macrophages (Fig. 1E-F).

### *S.* Typhi T3SS-dependent effectors contribute to intramacrophage replication

We next interrogated which T3SS effectors contribute to *S.* Typhi replication in human macrophages. To this end, we constructed 25 mutant strains that are deficient for previously identified T3SS-effectors and putative pseudogenes in *S*. Typhi (8). We then tested these mutants for replication defects in human macrophages using our time-lapse florescence dilution assay in 96-well dishes to facilitate higher throughput. Each dish contained wells infected with the WT *S*. Typhi strain and the Δ*phoP* and Δ*invA*Δ*ssaV S*. Typhi mutant strains which do not replicate in THP-1 macrophages. As an additional control we included an *S*. Typhi Δ*sptP* mutant, which lacks an effector gene that has been reported to be non-functional in *S.* Typhi (21). Consistent with these previous findings, we did not see a contribution of SptP to *S*. Typhi replication in macrophages (Figs. 2A, S2A).

**Figure 2.**
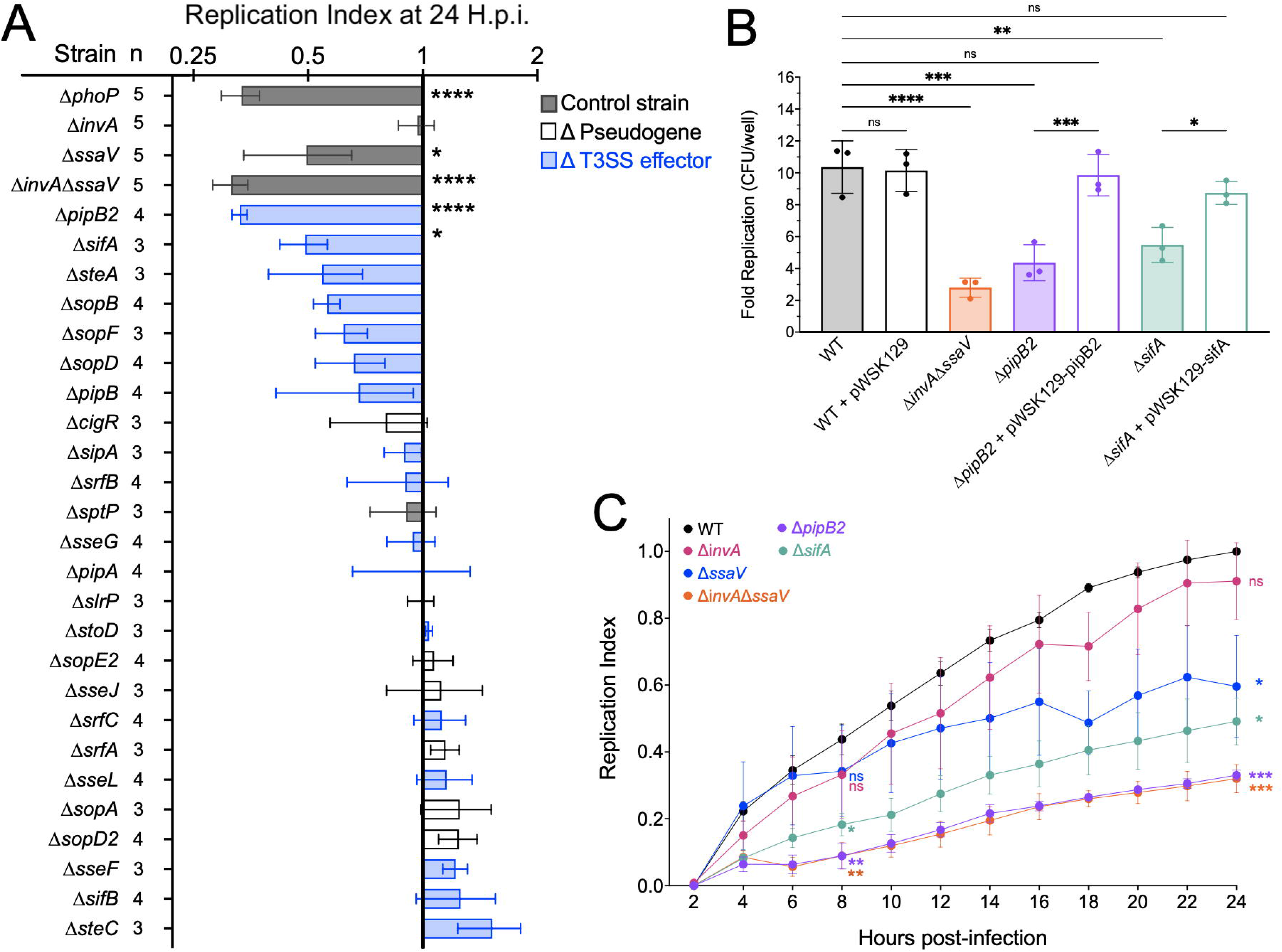
T3SS-dependent effectors contribute to intracellular replication. **A.** Replication index of each *S.* Typhi strain in THP-1 macrophages at 24 hours post-infection. T3SS effectors in ascending order of replication at 24 h.p.i. Statistical significance compared to hypothetical mean of 1.0 by Wilcoxon test. Each bar: mean of 3-5 biological replicates, each an average of 3 technical replicates. N = number of biological replicates performed for each strain. Colored by category of the gene knocked-out. Grey = control strain, expected outcome, Blue = T3SS effector knockout, White = T3SS effectors pseudogenized in *S.* Typhi. Error: SEM of biological replicates. **B.** Replication of *S.* Typhi in THP-1 macrophages by CFU/well at 2 and 24 h.p.i. Statistical significance by ANOVA. Dots: biological replicates, each an average of 3 technical replicates. Bars: mean. Bars filled with color indicate a disrupted gene, whereas bars filled in with white indicate the corresponding gene has been complimented. Error: SD. For all; ns or blank = p-value > 0.05, *≤0.05, ** ≤0.01, *** ≤0.001, **** ≤0.0001 **C.** Replication of *S.* Typhi in hMDMs measured by fluorescence dilution (pFCcGi) at throughout infection by time-lapse microscopy. Statistical significance compared to WT at 8 and 24 h.p.i. by ANOVA. Dots: mean of 3-5 biological replicates. Error: SEM. For all; ns = p-value > 0.05, *≤0.05, ** ≤0.01, *** ≤0.001, **** ≤0.0001

The results of this screen demonstrated that PipB2 and SifA contributed significantly to *S*. Typhi replication in human macrophages (Figs. 2A, S2A). We confirmed an intramacrophage replication defect for the *S*. Typhi Δ*pipB2* and Δ*sifA* mutant strains by plating viable bacteria (Fig. 2B). Importantly, the defects of the *S*. Typhi Δ*pipB2* and Δ*sifA* mutant strains was rescued by providing wild type copies of *pipB2* and *sifA*, respectively (Fig. 2B). In addition, we noticed that *S*. Typhi strains lacking SteA and SopB have slight defects in the fluorescence dilution assay (Fig. 2A), suggesting that they may contribute to intramacrophage replication, which is in agreement with previously published findings using *S.* Typhimurium models (22–26). However, we recovered the same CFUs for Δ*steA*, and Δ*sopB* mutant *S*. Typhi strains as the WT *S*. Typhi strain (Fig. S2B). Collectively, our results indicate that replication of *S*. Typhi in human macrophages is dependent on both PipB2 and SifA. The *S.* Typhi Δ*sifA* mutant as more severely attenuated than the T3SS-2 mutant (Δ*ssaV*) at 8 h.p.i. (Fig. 2C) with kinetics of intracellular replication similar to the Δ*pipB2* mutant (Fig. 2C). Previous studies with S. Typhimurium infection of murine macrophages have shown that PipB2 is translocated through T3SS-1 at early timepoints, then by T3SS-1 and -2 at later timepoint (27). Therefore, we hypothesized that the early translocation of SifA by *S*. Typhi is dependent on T3SS-1.

### *S.* Typhi Translocates PipB2 and SifA into macrophages through both T3SS-1 and -2

To assess translocation of *S.* Typhi T3SS effectors into the macrophage cytosol we constructed translational fusions between *S.* Typhi SifA and PipB2 with the TEM-1 β-lactamase reporter. GST fused to TEM-1 β-lactamase was used as a negative control. Translocation was detected in THP-1 macrophages using the fluorescent β-lactamase substrate CCF4-AM as described previously (28, 29). A fraction of macrophages infected with WT *S.* Typhi containing PipB2-BlaM or SifA-BlaM emitted blue fluorescence at 450 nm, whereas a GST-BlaM control did not, suggesting that fusions between these effectors and TEM-1 are translocated into host macrophages (Fig 3A). To determine whether translocation of SifA and PipB2 is dependent on T3SS-1 or -2, we measured translocation of SifA-BlaM or PipB2-BlaM into THP-1 macrophages infected with T3SS-1, -2 and T3SS-null *S.* Typhi strains at 8 and 16 h.p.i. (27). PipB2-BlaM translocation was reduced when either T3SS-1 or -2 was disabled at both timepoints (Fig 3B-C). Surprisingly, SifA-BlaM translocation was consistently reduced in the T3SS-1 mutant at both 8 and 16 h.p.i.. In contrast, the T3SS-2 mutant did not have a translocation defect at 8 h.p.i., but did have a translocation defect by 16 h.p.i. (Fig. 3B-C). We further confirmed T3SS-1 dependent translocation of SifA-BlaM in primary human macrophages by confocal microscopy of hMDMs infected with WT, Δ*invA*, Δ*ssaV*, and Δ*invA*Δ*ssaV S*. Typhi strains (Fig. S3). Taken together, these results indicate that *S.* Typhi replication in human macrophages is dependent on the effectors PipB2 and SifA and that translocation of PipB2 and SifA into macrophages is dependent on the presence of either T3SS-1 or -2.

**Figure 3.**
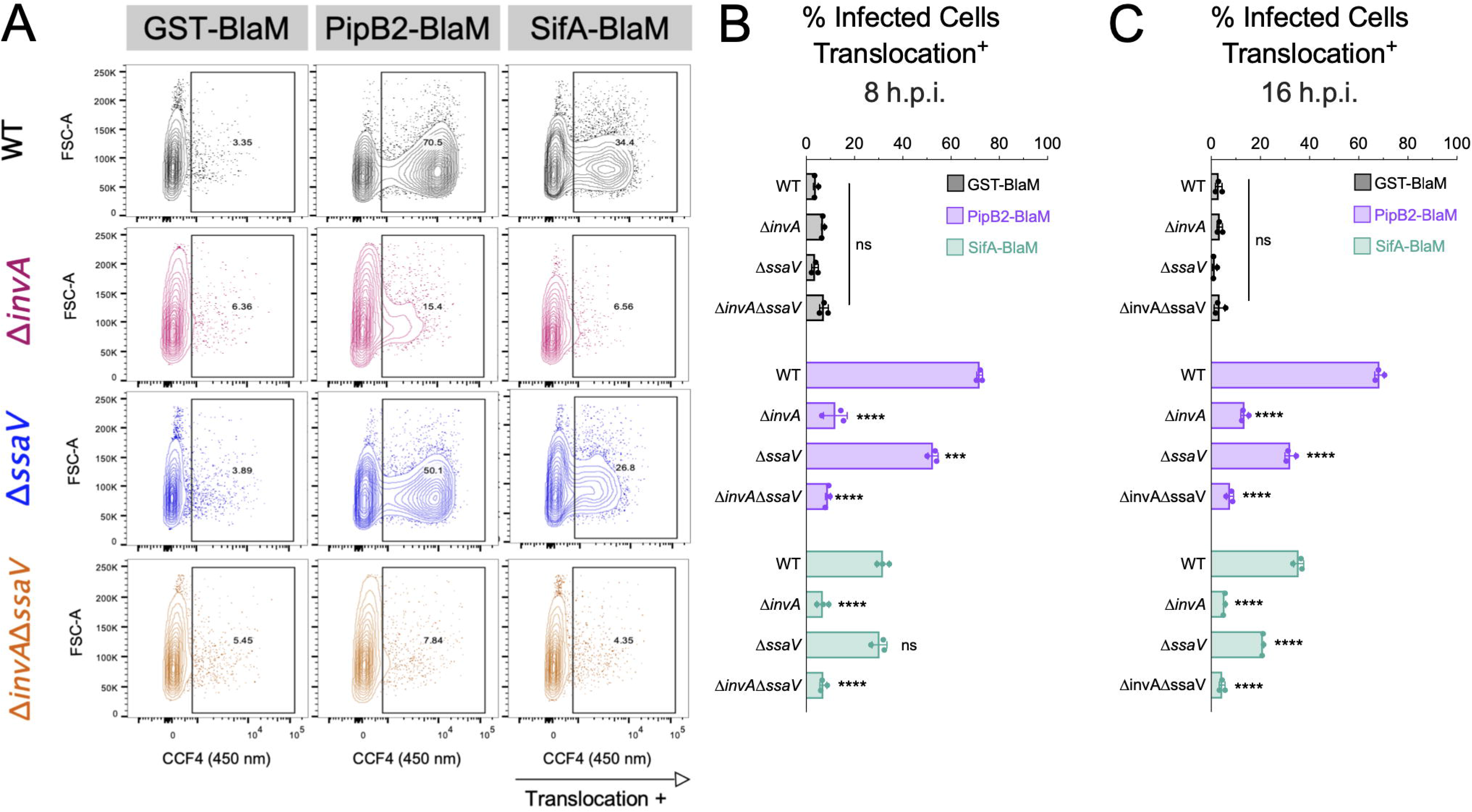
Translocation of SifA by 8 h.p.i. is T3SS-1-dependent, whereas translocation of SifA by 16 h.p.i. is dependent on both T3SS-1 and -2. **A.** Representative flow cytometric analysis of THP-1 macrophages infected with *S.* Typhi expressing effector-BlaM constructs at 8 h.p.i. Columns = GST-BlaM, PipB2-BlaM, SifA-BlaM translocation at 8 h.p.i. detection in hMDMs. Rows = strain of Ty2 used to infect, carrying BlaM construct. **B.** Quantification of tranlocation assay in samples fixed at 8 h.p.i., statistical significance by 2- way ANOVA. Dots: technical replicates. Bars: mean of technical replicates. Representive of 3 biological replicates. ns = p-value > 0.05, ***≤0.001, **** ≤0.0001 **C.** Quantification of tranlocation assay in samples fixed at 16 h.p.i., statistical significance by 2- way ANOVA. Dots: technical replicates. Bars: mean of technical replicates. Representive of 3 biological replicates. ns = p-value > 0.05, **** ≤0.0001

### *S.* Typhi T3SS-1 and T3SS-2 contribute to virulence in a humanized mouse model of typhoid fever

Although *S.* Typhi is human-restricted, researchers have described systemic infection in mice with “humanized” immune systems (30). A recent study aimed at identifying *S.* Typhi virulence factors required for acute infection in NOD-*Prkdc^scid^IL2rg^tm1Wjl^*(NSG) mice engrafted with human CD34^+^ hematopoietic stem cells derived from umbilical cord blood (hu-SRC-SCID mice) identified Vi capsule, lipopolysaccharide (LPS), aromatic amino acid biosynthesis and the siderophore salmochelin as essential for virulence (13). However, they found that the T3SS-2 did not provide a competitive advantage during the first 48 h of infection of hu-SRC-SCID mice. Based on our *in vitro* findings in human macrophages, we hypothesized that both *S.* Typhi T3SSs contribute to virulence in the hu-SRC-SCID model. To test this idea, hu-SRC-SCID mice were infected as described by Karlinsey et al. Briefly, hu-SRC-SCID mice were infected intraperitoneally (IP) with an equal mixture of *S.* Typhi WT and isogenic Δ*invA*Δ*ssaV* mutant strain (10^5^ CFU of each). The competitive index (CI) was calculated for the spleen and liver at 2 days p.i. The mutant lacking both T3SSs was not outcompeted by the WT strain (Fig. 4A). However, we reasoned that a later time point may reveal a role for the T3SSs because the bacteria would have more time to replicate in human-derived macrophages in the humanized mice. To test this idea, we infected the hu-SRC-SCID mice IP with a 10-fold lower dose that contained an equal mixture of two strains (10^4^ CFU of each). Mice were infected with *S.* Typhi WT and isogenic mutants: either *sptP::kan^R^* as a control for kanamycin resistance inserted into a non-functional gene (21), an Δ*ssaV::kan^R^* strain or an Δ*invA*Δ*ssaV::kan^R^* strain. Strikingly, at 5 days p.i. the T3SS-null *S.* Typhi strain was significantly outcompeted by the WT strain in both the spleens and the livers (Fig. 4B). In contrast, the Δ*sptP* mutant *S.* Typhi strain was not outcompeted by the WT strain in the spleen or liver (Fig. 4B). Although deletion of T3SS-2 alone had a defect relative to the control strain in the spleen, it was not significantly outcompeted by WT in the liver, and it was not out-competed as significantly as the T3SS-null mutant in either organ. Previous publications using this humanized mouse model of typhoid infection have highlighted the heterogeneity in the levels of *S*. Typhi recovered from individual mice (13, 30). However, we recovered greater than 10^3^ WT *S.* Typhi CFUs per gram of tissue in 11 of the 12 spleens (Fig. S4A), indicating that the spleens contained an amount of WT bacteria well above the limit of detection. Liver burdens were more heterogenous than spleen burdens (Fig. S4B), however the level of WT bacteria were 10-fold above the limit of detection, at 10^2^ CFU per gram of tissue. Together, these data demonstrate that *S.* Typhi uses two T3SSs to colonize systemic sites in a humanized mouse model of typhoid fever.

**Figure 4.**
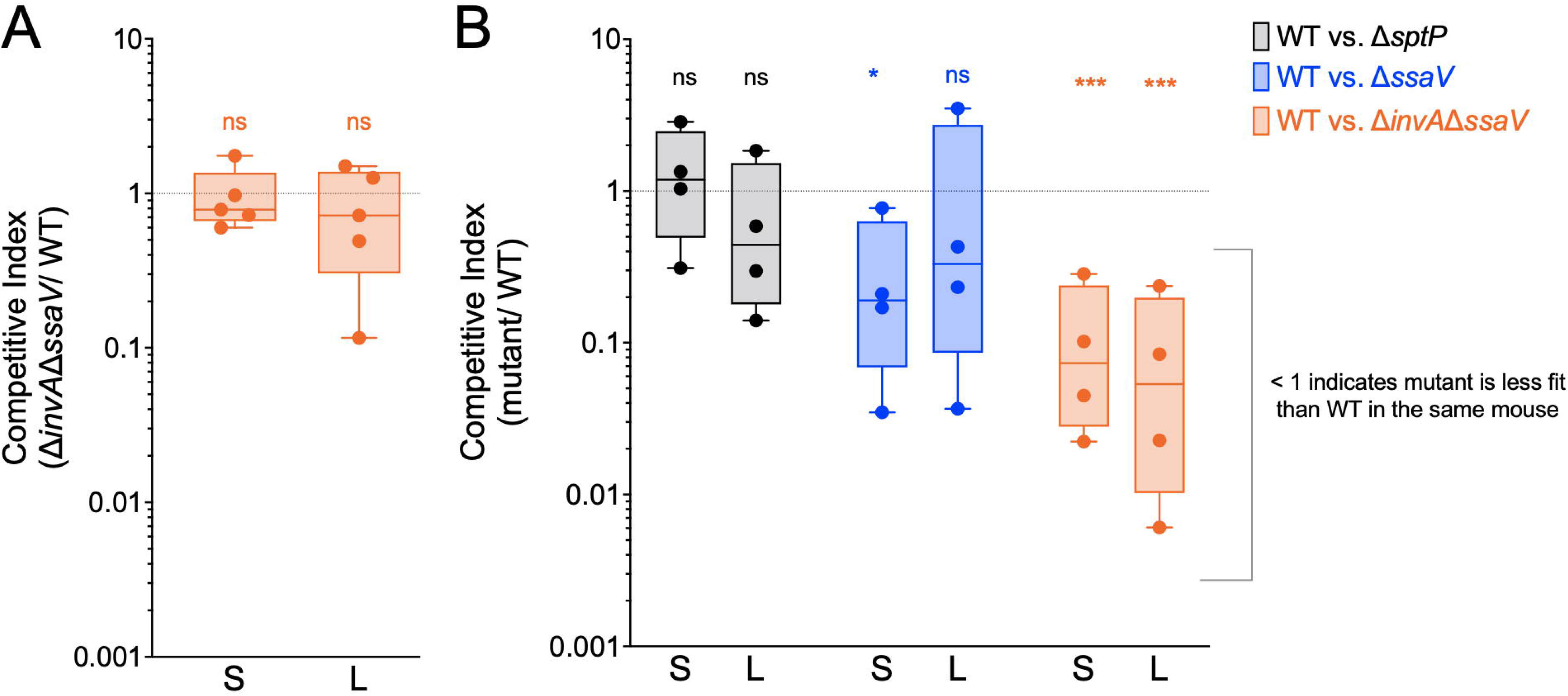
*S.* Typhi T3SS-1 and -2 contribute to virulence in a humanized mouse model of typhoid fever. **A.** Humanized mice infected with 2x10^5^ bacteria intraperitoneally. Competitive index of viable bacteria recovered from spleen and liver 2 days p.i.. Statistical significance by one-Sample t-test compared to a theoretical mean of 1.0. S= spleens. L: livers. Dots: individual mice. ns = p-value > 0.05 **B.** Humanized mice infected with 2x10^4^ bacteria intraperitoneally. Competitive index of viable bacteria recovered from spleen and liver 5 days p.i.. Statistical significance by One-Sample t-test compared to a theoretical mean of 1.0. S: spleens. L: livers. Dots: individual mice. ns = p-value > 0.05, * ≤0.05, *** ≤0.001

## Discussion

Many studies have compared the genomes of the host generalist *S.* Typhimurium with *S.* Typhi and have speculated that the evolution of *S.* Typhi likely involved both gain of function and loss of function as it evolved to selectively infect humans (8). Although systemic murine infection with *S.* Typhimurium models human disease, fundamental genetic differences between typhoidal and non-typhoidal *Salmonella* serovars limits the ability of this model to identify human-specific virulence mechanisms. In this study, we used human-derived macrophages and a humanized mouse model to study *S*. Typhi virulence factors that contribute to intramacrophage replication (13, 30). We show that both T3SS-1 and -2 promote *S.* Typhi replication in human macrophages, and further that both T3SSs contribute to *S.* Typhi colonization of the spleen and liver in a humanized mouse model of typhoid fever. There is clinical interest in determining if T3SS-null *S.* Typhi may be useful as an attenuated vaccine strain (31). This study provides evidence that *S.* Typhi mutants lacking both T3SS-1 and -2 are attenuated for intramacrophage replication. A previously published clinical study of an *S.* Typhi Δ*aroA*Δ*ssaV* mutant, which lacks a functional T3SS-2, was administered to human volunteers, with infrequent adverse events, indicating that *S.* Typhi Δ*aroA*Δ*ssaV* is likely attenuated within the specific patient population studied (31). The data presented here show T3SS-1 and -2 deletion attenuates the laboratory *S.* Typhi strain Ty2, which is closely related to the only currently licensed live, attenuated typhoid vaccine strain Ty21a (32). However, more rigorous clinical studies appropriately accounting for the diversity in genetics and lifestyle of all effected populations are necessary (3).

Previous research has demonstrated that while T3SS-2 is required for *S.* Typhimurium replication in murine macrophages, T3SS-2 is not required for *S.* Typhi replication in human macrophages (12, 33). Here, we used a fluorescence dilution assay that directly assesses bacterial replication in human macrophages (Fig. 1B-C). We uncovered that T3SS-2 partially contributes to *S.* Typhi replication in human macrophages. We further showed that disabling both T3SS-1 and -2 renders *S.* Typhi unable to replicate in human macrophages.

To gain further insights into the potential roles of *S*. Typhi T3SS effectors we screened 25 strains lacking individual T3SS effectors for replication defects in which mutant strains were simultaneously analyzed in a time-lapse fluorescence dilution assay. Many of the single mutants did not have significant replication defects, which would be expected for the reported pseudogenes (*sptP*, *cigR*, *sopA*, *srfA*, *sopE2*, *slrP*, *sopD2*, and *sseJ*). We also found that the *S*. Typhi Δ*steC* mutant had the highest replication index at 24 h.p.i., although it didn’t quite reach statistical significance (Fig. 2A). This finding agrees with previously published data indicating SteC may function to restrain intracellular growth during *S.* Typhimurium infection (34). Future studies are warranted to examine typhoidal SteC function in human macrophages specifically.

Our screen also identified two T3SS effectors, PipB2 and SifA, that contribute significantly to replication in human macrophages (Fig. 3A-C). We confirmed that the effector proteins PipB2 and SifA are important for intramacrophage growth of *S*. Typhi by plating for viable bacteria (Fig. 2B). We also show that both T3SS-1 and -2 contribute to translocation of these important effectors into the cytosol of human macrophages and that SifA translocation is T3SS-1-dependent at 8 h and that T3SS-2 contributes to translocation at 16 h.p.i. (Fig. 3B-C). Previous studies in *S*. Typhimurium have shown that PipB2 is translocated through both T3SS-1 and -2 (27). Similarly, our results indicate that PipB2 can be translocated by *S*. Typhi’s T3SS-1 and -2 (Fig. 3B-C). In contrast, others have shown that SifA is translocated through T3SS-2 in *S*. Typhimurium (35). Interestingly, our results show that in *S.* Typhi SifA can be translocated into the macrophage cytosol by T3SS-1 at 8 h.p.i. and both T3SS-1 and -2 at 16 h.p.i. (Fig. 3B-C). Consistent with this finding, there are fewer replicating *S*. Typhi Δ*sifA* mutant bacteria at 8 h.p.i. compared to the *S*. Typhi T3SS-2 Δ*ssaV* strain (Fig. 2C). Finally, the kinetics of intramacrophage replication of the *S*. Typhi Δ*sifA* and Δ*pipB2* mutants were similar, indicating that these effectors play important roles in establishing and maintaining the SCV during *S*. Typhi infections of human macrophages (Fig. 2C). In *S*. Typhimurium PipB2 and SifA are involved in SCV membrane dynamics (36). For example, both PipB2 and SifA recruit kinesin-1 to the SCV through a direct interaction or via SKIP, respectively (37–39). SifA also recruits Rab9 and is thought to suppress lysosome functions (40, 41). Future studies focused on effector-host protein- protein interactions during *S*. Typhi infections of human macrophages will be important for increased knowledge of host-pathogen interactions during typhoid fever. Finally, Figueira et al. demonstrated in *S*. Typhimurium that poly-effector mutant strains are more severely attenuated in replication compared to single mutant strains, suggesting some redundancy in effector functions (11). For example, the effectors SseF and SseG directly interact and cooperate during S. Typhimurium infection (25, 42, 43). This may partially explain why *S.* Typhi Δ*sseF* and Δ*sseG* mutant strains did not have significant intramacrophage replication defects. However, these two effectors each individually significantly contribute to *S.* Typhimurium intramacrophage replication (11). To probe specific differences in effector functions between serovars, a direct comparison of serovar replication within the same host cell context, including single and double-effector knockouts, and heterologous expression of homologous effectors in different serovars, will be necessary. For example, a thorough study of the genetic and molecular reasons for SptP functional differences between *S.* Typhi and *S.* Typhimurium homologs revealed that typhoidal SptP loss-of-function is caused by a mutation in the chaperone-binding domain (21). It is possible that other typhoidal *Salmonella* effectors also have mutations that cause either divergent functionality or loss-of-function compared to their S. Typhimurium homolog. On the other hand, *S.* Typhi has a significantly reduced T3SS effector repetoire. For example, SopD2, which is pseudogenized in *S.* Typhi, modulates trafficking of the SCV and has recently been shown to cooperate with SteA and PipB2 (11, 25, 44, 45). Therefore, a possible explanation for why some individual effectors, such as PipB2, play a more significant role for intramacrophage replication of *S.* Typhi compared to S. Typhimurium, is that the functionally redundant effectors are missing in *S*. Typhi (11). Future studies including analysis of poly- effector mutants of *S*. Typhi will be important to increase our understanding of T3SS effector functional redundancies and possibly explain why PipB2 significantly contributes to *S*. Typhi intramacrophage replication.

Here we show that *S*. Typhi intramacrophage replication is dependent on T3SSs activity. However, previous studies have shown that T3SSs also translocate flagellin and structural components of the T3SS into the cytosol which are pathogen-associated molecular patterns (PAMPS) that are recognized by pattern recognition receptors (PRRs) (18, 19, 29, 46). Indeed, human macrophages produce a cytosolic PRR known as hNAIP (the NLR [nucleotide-binding domain, leucine-rich repeat-containing] family, apoptosis inhibitory protein) that recognizes flagellin (19, 46) and the T3SS-1 inner rod protein and the T3SS-1 and -2 needle proteins (18), triggering a signaling cascade that results in a pyroptotic cell death. However, we do not see high levels of cell death in THP-1 macrophages when infected with WT or mutant *S.* Typhi strains in this study (Fig. S1D). Future studies are therefore warranted to examine host cell detection of intramacrophage *S.* Typhi replication and whether *S*. Typhi has mechanisms to effectively evade detection by the macrophage.

Importantly, we demonstrated the pathophysiological relevance of our *in vitro* studies by showing that the T3SSs are critical for *S*. Typhi colonization in systemic tissues of humanized mice. Functional redundancy in *S.* Typhi’s T3SSs likely explain why previous screens for *S.* Typhi genes required for intramacrophage survival and systemic colonization of hu-SRC-SCID mice failed to identify T3SS-1 and -2 as crucial virulence factors (13, 47). Overall, the data presented here demonstrate that both T3SS-1 and -2 are critical virulence factors enabling *S.* Typhi replication in systemic tissues, and specifically within human macrophages.

## Materials and Methods

### Key Resources Table

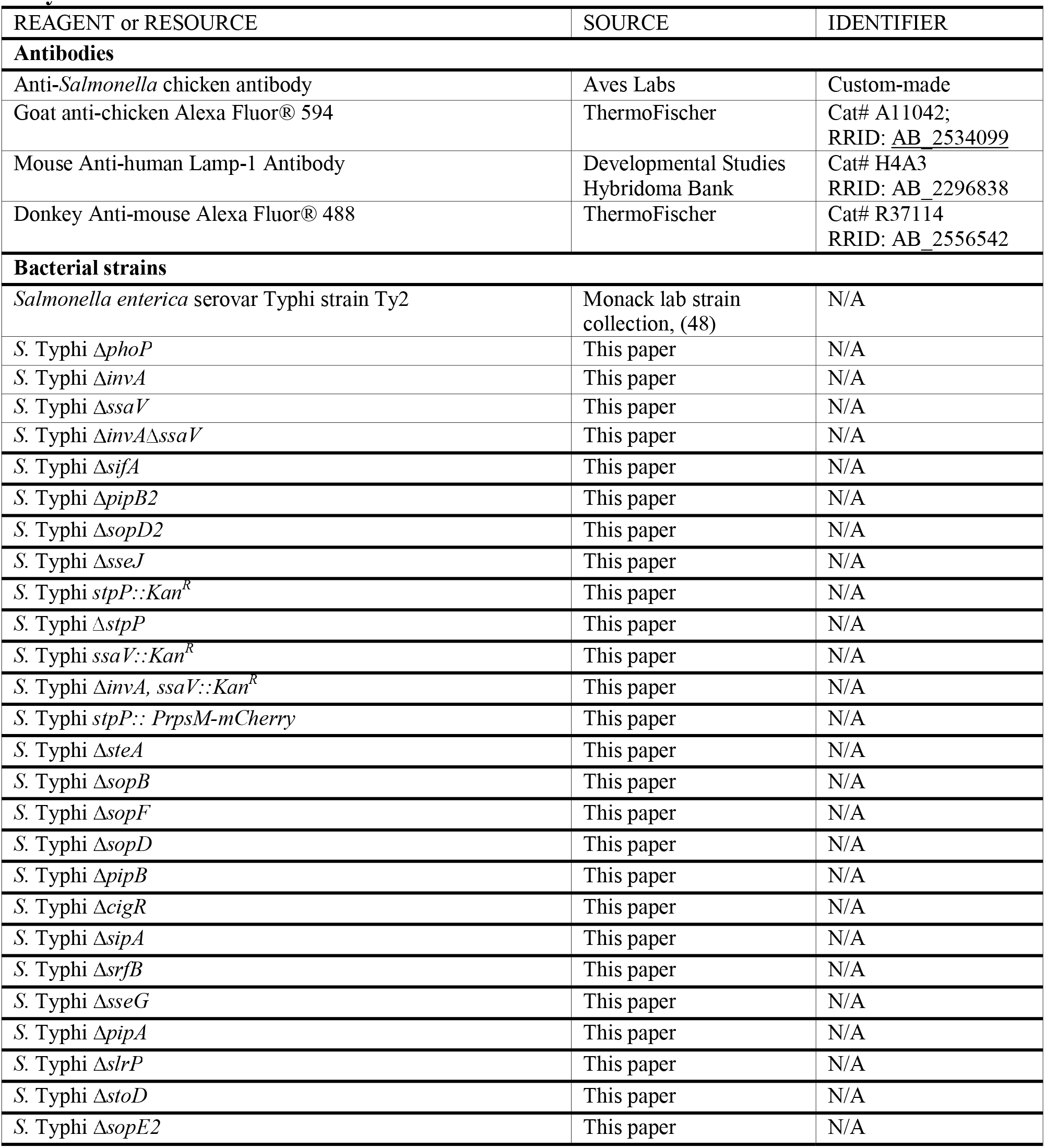
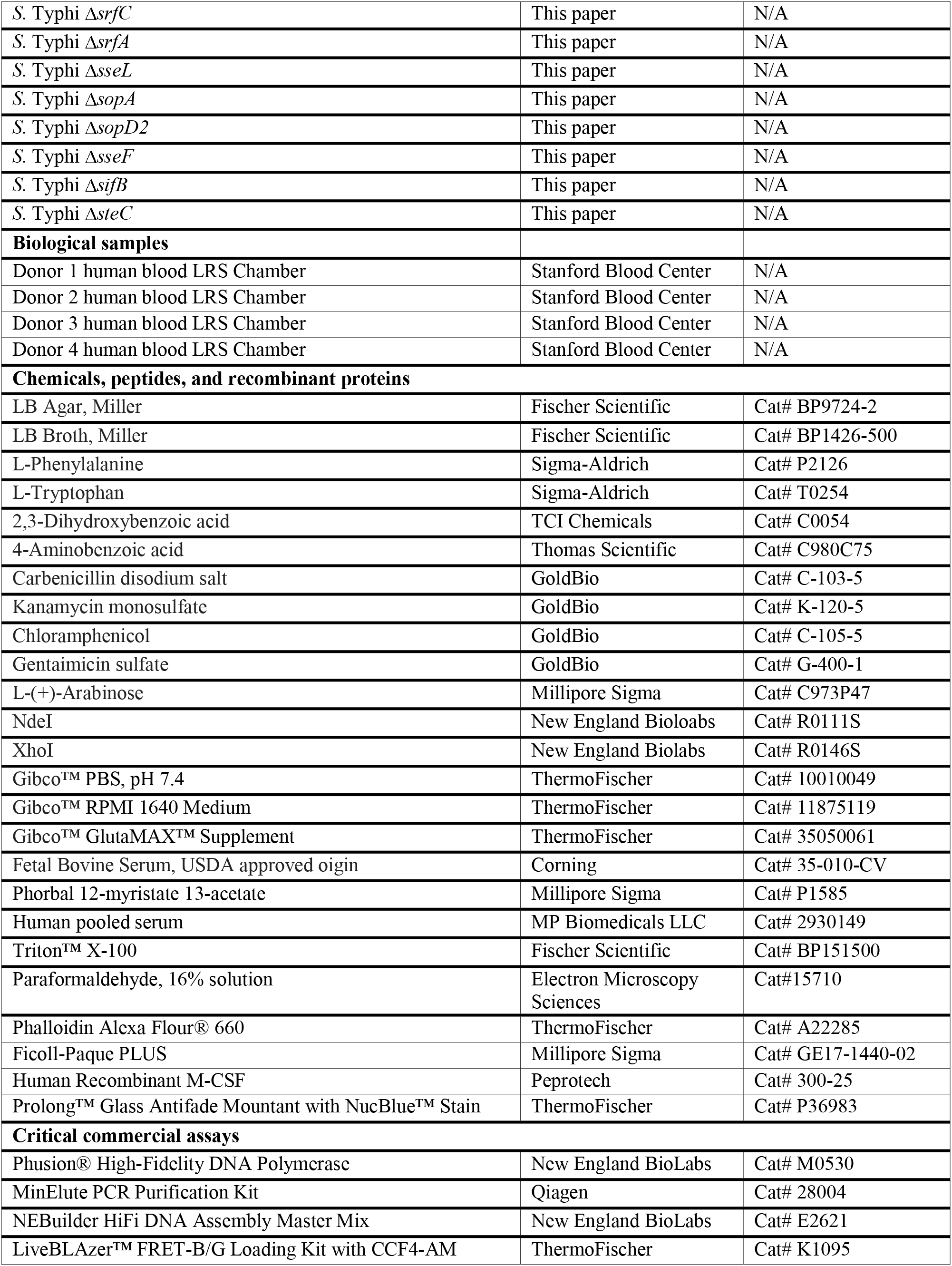
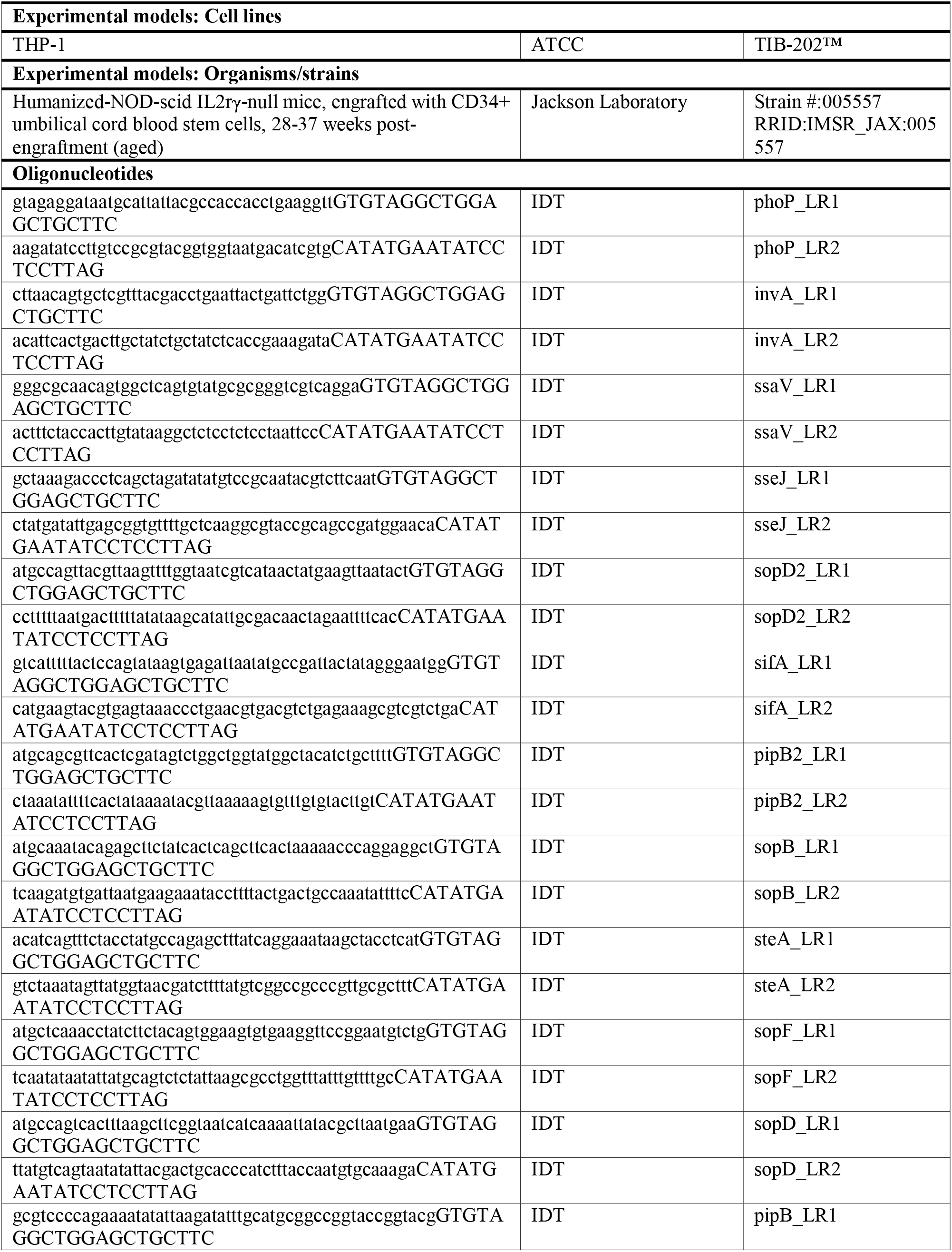
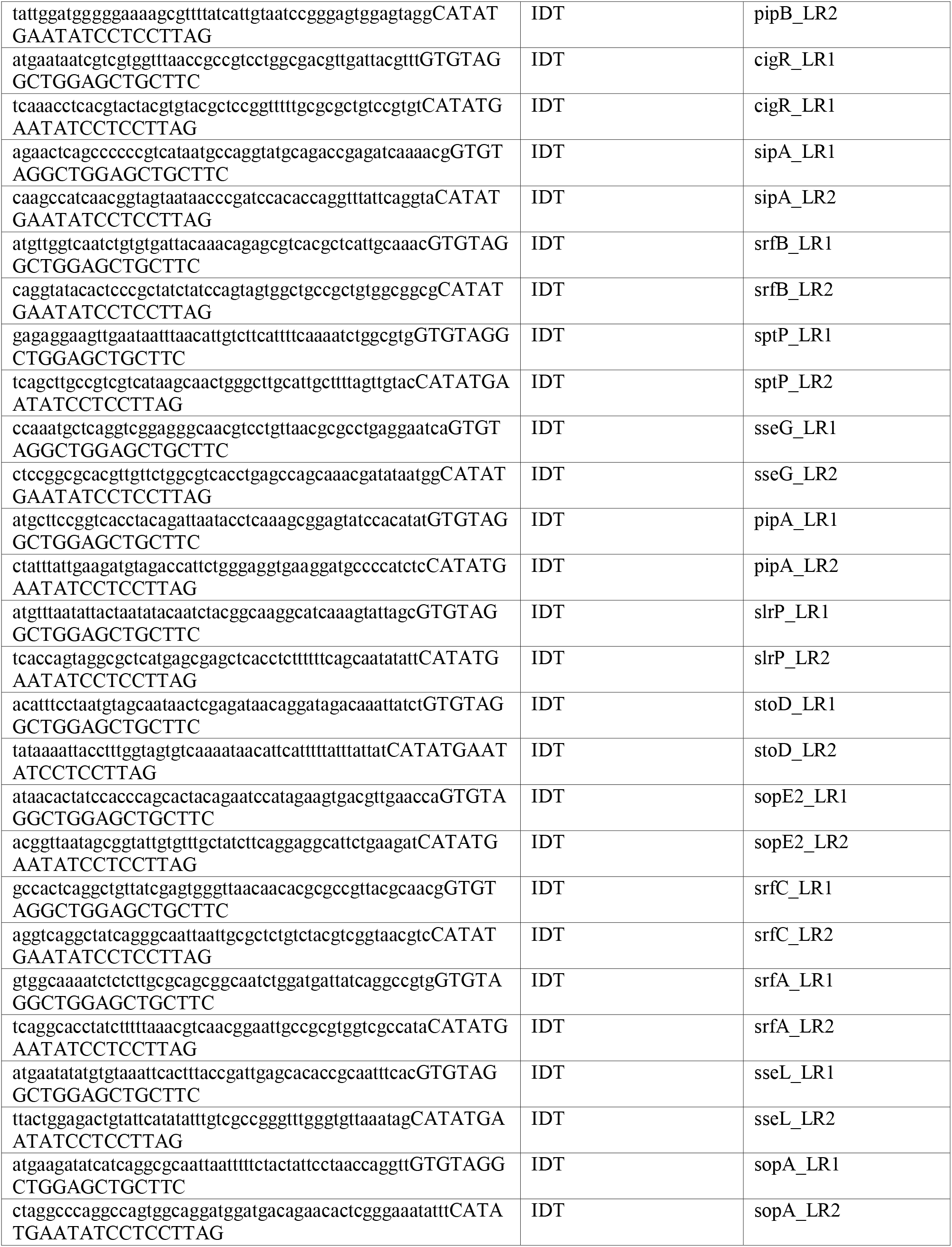
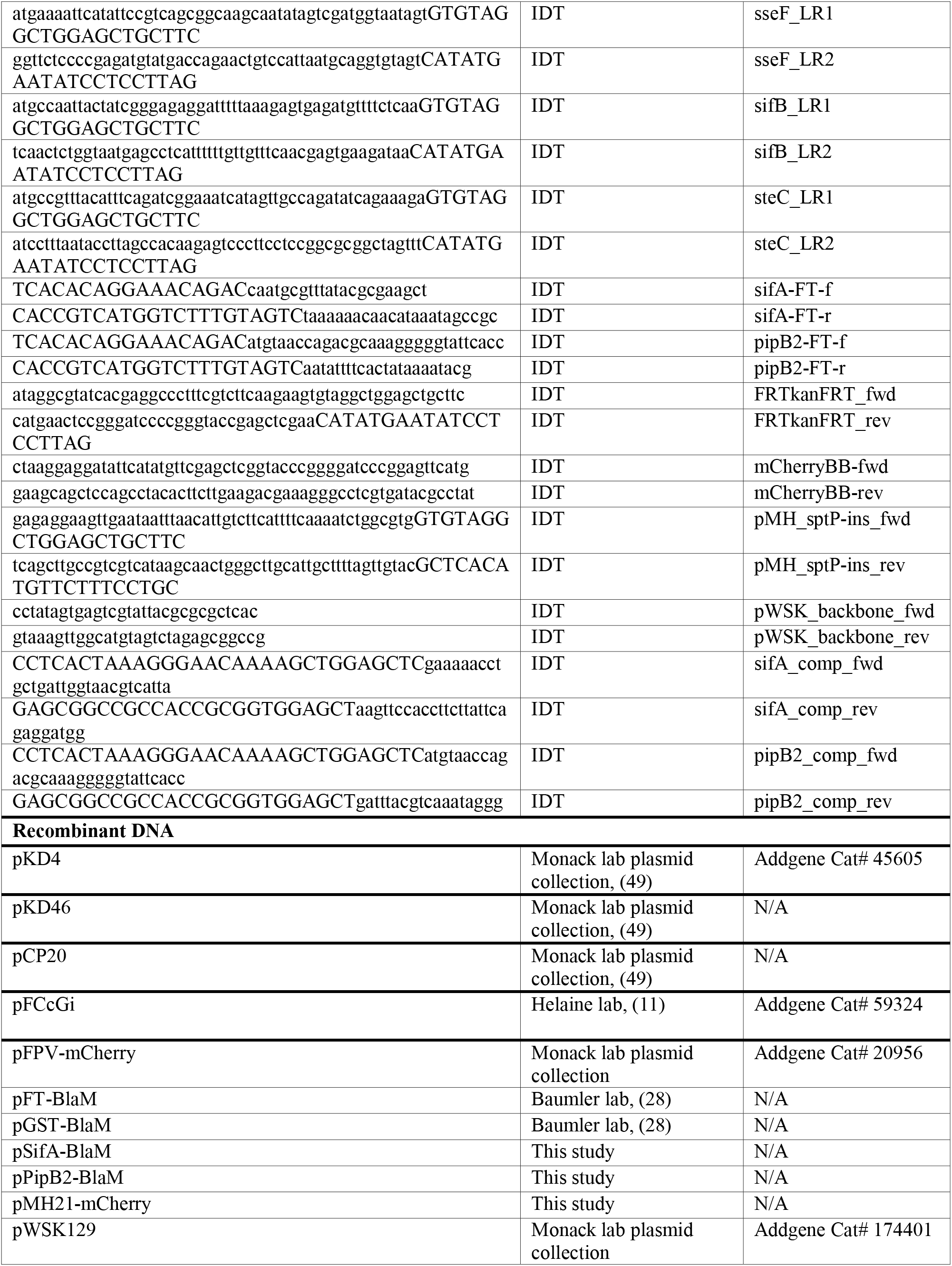
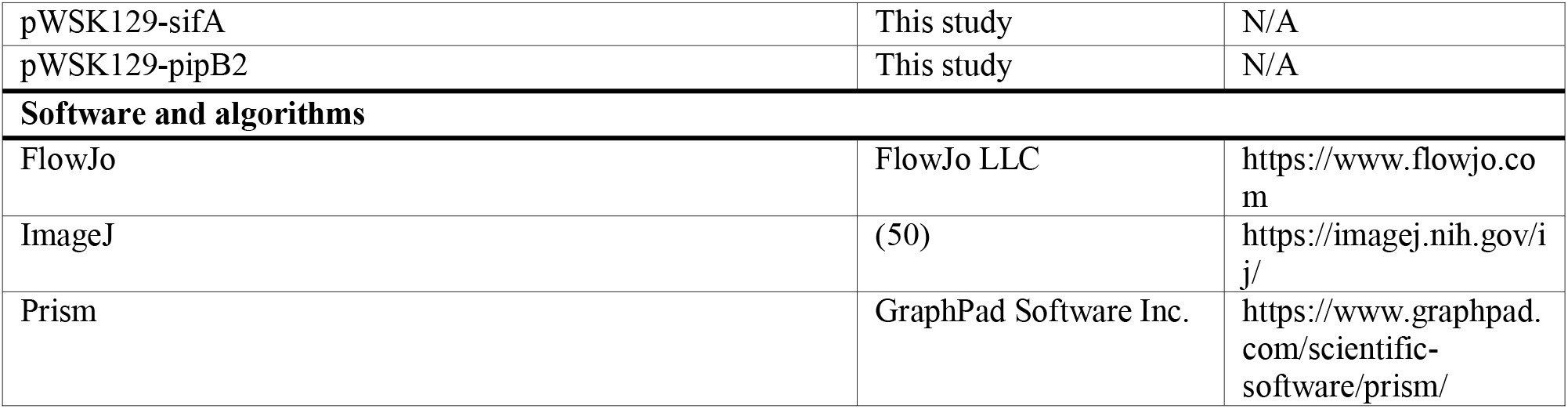

### Mouse Strains and Husbandry

Experiments involving animals were performed in accordance with NIH guidelines, the Animal Welfare Act, and US federal law. All animal experiments were approved by the Stanford University Administrative Panel on Laboratory Animal Care (APLAC) and overseen by the Institutional Animal Care and Use Committee (IACUC) under Protocol ID 12826. Animals were housed in a centralized research animal facility accredited by the Association of Assessment and Accreditation of Laboratory Animal Care (AAALAC) International. Female hu-SRC-SCID mice engrafted with CD34+ human umbilical cord blood stem cells were obtained from Jackson Laboratories. Humanized-NOD-scid IL2rγ-null mice, engrafted with CD34+ umbilical cord blood stem cells, 28-37 weeks post-engraftment (aged) (Jackson, Strain #:005557) were housed under specific pathogen-free conditions in filter-top cages that were changed weekly. Sterile water and food were provided ad libitum. Mice were given at least one week to acclimate prior to experimentation.

### Cell culture and differentiation

THP-1 cells were obtained from ATCC and passaged a maximum of 10 times in all experiments. THP-1s were routinely cultured in RPMI medium with 10% heat-inactivated FBS and 2 mM GlutaMAX supplement. THP-1 cells were differentiated into macrophage-like adherent cells according to published protocols (17). Briefly, THP-1s were treated with 100 nM PMA for 48 hours, then media was replaced and THP-1s were infected 3 days after differentiation. To better adhere differentiated THP-1s, before seeding plates for differentiation the surface of TC-treated plastic was coated with human plasma-derived fibronectin according to the manufacturer’s instructions.

For primary human-derived macrophages four individual TrimaAccel® LRS chambers recovered after Plateletpheresis containing white blood cell concentrate were purchased from Stanford Blood Center. Each chamber contained samples from an individual platlet donor. Samples were entirely de-identified, products collected and sold by Stanford Blood Center for *in-vitro* investigational use are not routinely tested for infectious disease markers and do not require IRB approval. Peripheral blood mononuclear cells were isolated by gradient centrifugation using Ficoll-Paque PLUS, and adherent mononuclear cells were differentiated with 30 ng/ml hM-CSF in RPMI medium supplemented with GlutaMAX and 10% heat-inactivated FBS for 6-7 days prior to infection as previously described (48).

### Bacterial Strains and Growth Conditions

*S.* Typhi was routinely cultured in LB broth supplemented with “aromix” (4 mg/mL L-phenylalanine, 4 mg/mL L-tryptophan, 1 mg/mL p-aminobenzoic acid, 1 mg/mL dihydroxybenzoic acid), and antibiotic selection when indicated at the following concentrations: kanamycin 50 µg/mL, carbenicillin 50 µg/mL, chloramphenicol 25 µg/mL.

### Bacterial Strain Construction

Genes in the Ty2 genome were targeted for deletion following an adjusted lambda red mutagenesis protocol (49). Briefly, primers provided in the key reagents table were used to amplify the kanamycin resistance cassette of pKD4, and PCR products were purified and concentrated to 500 ng/uL using a MinElute PCR product purification kit (Qiagen). Strains containing pKD46 were grown at 30 °C, 200 rpm until mid-log phase, then L-arabinose was added to a final concentration of 50 mM, and incubated for an additional 1-2 h. Bacteria were pelleted and washed in ice-cold autoclaved DI water, concentrated 200X, and 1-5 µg of purified PCR product was added to each electroporation cuvette. Bacteria were electroporated then recovered in SOC broth at 30 °C for 3 h static. Cultures were plated on antibiotic selection plates and grown overnight at 42 °C. Colonies were PCR verified for kanamycin resistance insertion and endogenous gene disruption. Then pCP20 was electroporated into kanamycin resistant strains, and removal of the kanamycin resistance cassette was PCR confirmed. Removal of both the pKD46 and pCP20 plasmids was confirmed by a loss of carbenicillin resistance. pFCcGi was electroporated into strains and selected for with 50 µg/mL of carbenicillin. When indicated, 0.4% L-arabinose was added to pFCcGi+ cultures to induce GFP expression.

For chromosomal insertion of constitutive mCherry, the plasmid pMH21-mCherry was first constructed. The kanamycin resistance cassette from pKD4 was amplified using FRTkanFRT_fwd and FRTkanFRT_rev and the backbone of pFPV-mCherry, designed for constitutive mCherry expression in *Salmonella* was amplified using mCherryBB-fwd and mCherryBB-rev. Linear PCR products were digested with DpnI, purified, and ligated with NEBuilder HiFi DNA Assembly Master Mix according to the manufacturer’s instructions to create pMH21-mCherry. Then, pMH_sptP-ins_fwd and pMH_sptP-ins_rev were used to amplify the kanamycin resistance cassette, the PrpsM promoter, and mCherry gene from pMH21- mCherry, adding homology with the sptP locus. Mutagenesis proceeded as previously stated using the lambda red method, including using pCP20 to then remove the kanamycin resistance cassette. Each bacterium thus encodes one constitutive mCherry locus in place of the *sptP* gene.

### Beta-Lactamase Translocation Constructs

The pFT-BlaM vector and pGST-BlaM were provided by the Baumler lab (28). To construct the sifA-FT and pipB2-FT plasmids, the pFLAG-TEM1 vector was digested with *Nde*I and *Xho*I and fragments of Ty2 gDNA were amplified using primers listed in the Key Resources Table. All purified plasmids were sequenced by Plasmidsaurus (Eugene, Oregon). Plasmids were electroporated into Ty2 *sptP*::*mCherry*.

### THP-1 Infections

Bacteria were grown in LB+Aromix broth with 0.4% arabinose at 25 °C, 200 rpm (Brewer et al., 2021) until reaching an OD of ∼1.0. Bacterial pellets were spun down at 3500 rpm then washed and diluted with PBS to the appropriate CFU/mL to reach a final multiplicity of infection of 1:20 and opsonized with 25% pooled human serum for 30 minutes at room temperature. Bacteria were then resuspended using a sterile 25G needle and into the infection medium (RPMI + 10% heat inactivated FBS + 2 mM GlutaMAX). To synchronize uptake, plates were spun at 250 xG for 5 minutes, then incubated at 37 °C, 5% CO2 for 1 hour. Wells were washed with RPMI then incubated in infection medium with 100 µg/mL gentamicin for 1 hour to kill extracellular bacteria. Wells were then washed and incubated in infection medium containing 20 µg/mL of gentamicin for the remainder of the infection. When indicated, 2 h.p.i. corresponds to samples collected after the final wash.

### hMDM Infections

hMDM infections were performed similar to THP-1infections, except infections were performed with an MOI of 10:1 and for 30 minutes, followed by 1.5 hours of 100 ug/mL gent treatment.

### Flow Cytometry Analysis of Intramacrophage Replication

At indicated timepoints, intramacrophage replication of S. Typhi was quantified by flow cytometry according to previously published protocols (11, 20). Briefly, macrophages were lysed by incubation with 1% Triton-X at 37 °C for 10 minutes. Samples were pelleted at 13,000 xG for one minute then fixed in 4% paraformaldehyde for 15 minutes at room temperature. Samples were then washed with PBS once and resuspended in FACS buffer. 100,000 events were collected per sample on a BD LSRII flow cytometer and analyzed using FloJo software. Bacteria were gated on size relative to buffer-only control, and mCherry+ signal relative to non-fluorescent bacteria. The geometric mean of the GFP signal for mCherry+ events was then calculated for each sample. Technical replicates correspond separate wells infected at the same time, biological replicates correspond to infections performed on separate days.

### Quantifying CFU/well from Infected Macrophages

At indicated timepoints macrophages were lysed with 1% Triton-X at 37 °C for 10 minutes. Samples were pelleted at 13,000 xG for one minute and media was aspirated until 100 µL remained. Samples were resuspended, serially diluted in PBS and plated on LB + aromix plates. After overnight incubation at 30 °C, colony forming units (CFUs) were counted.

### Time-Lapse Microscopy

An Incucyte® S3 Live-Cell Analysis Instrument was used for time-lapse imaging. At indicated time intervals, Incucyte® Base Software was used to collect 20X images at a set location, 3-4 images per well were routinely collected. To quantify fluorescence, the Incucyte® Cell-By-Cell Analysis Software Module segmented all images into extracellular and intracellular area. To indicate replication, the ratio of intracellular mCherry intensity was divided by the intracellular GFP integrated intensity and plotted over time. For each biological replicate, three wells, 3-4 images each well per timepoint, were averaged together to indicate one replicate. Biological replicates indicate experiments performed on separate days. To compare biological replicates, values from uninfected wells were subtracted, then values were normalized to make the greatest value the WT culture achieved in each experiment equal to one and the lowest value collected in each experiment equal to zero, this value is named “Replication Index” (11). Individual images from the incucyte software were exported and compiled into videos using ImageJ.

### Optical Density Growth Curves

Cultures were grown in LB + aromix overnight at 37 °C, 200 rpm shaking, then diluted 1:50 and grown in LB + aromix for 3 hours to reach log phase. Aliquots of the cultures were then pelleted and washed three times in PBS, then optical density (OD600) was measured, and strains were back-diluted to a starting OD of 0.02. in 96-well plates. Plates were incubated at 37 °C with shaking between reads on a Synergy HTX (BioTek) plate reader and optical density was measured every 15 minutes. Biological replicates indicate experiments performed on separate days.

### Confocal Microscopy

Cover glass was coated with human fibronectin according to the manufacturer’s instructions. Macrophages were seeded, differentiated and infected on fibronectin-coated glass. At indicated time points, media was removed and infections were fixed with PLP fixative (2% paraformaldehyde in 75 mM NaPO4 buffer pH 7.4, 2.5 mM NaCl) for 15 mins at room temperature. Slides were gently washed after fixing and between stains with warm PBS supplemented with 9 mM CaCl2 and 5 mM MgCl2. After permeabilization with 1% saponin and 3% BSA, slides were first stained with 1:200 Mouse monoclonal LAMP-1 (DSHB) and 1:1000 Chicken anti-*Salmonella* (Aves Labs) for 1 hour at room temperature, followed by incubation in secondary antibodies 1:500 goat anti-chicken Alexa594 (Invitrogen), 1:500 donkey anti-mouse Alexa488 (ThermoFischer), and 1:100 Alexa660 Phalloidin (ThermoFischer) for 1 hour at room temperature. Cover glass was then mounted on slides with ProLong Glass Antifade with NucBlue stain (ThermoFischer). Slides were cured for 24 hours in the dark at room temperature and stored at -20°C until imaging on a Zeiss LSM 700 confocal microscope with the ZEN 2010 software. Images were processed using ImageJ. For imaging of beta-lactamase transloation into hMDMs, cells were seeded and infected on poly-L-lysine coated glass wells. At 8 h.p.i. cells were loaded with CCF4-AM dye according to the manufacturer’s instructions for 1.5 hours in the dark. Cells were then fixed with 3.2% PFA for 20 minutes in the dark. Cells were then washed once with PBS and PBS was used to keep the cells hydrated during imaging. Cells were imaged a Zeiss LSM 700 confocal microscope with the ZEN 2010 software with a blue diode 405 nm laser for excitation and with detection filters set at 410–450 nm for coumarin and 493–550 nm for fluorescein.

### Flow Cytometry Analysis of Effector Translocation

At 8 h.p.i., THP-1 macrophages were acclimated to room temperature and loaded with CCF4- AM dye according to the manufacturer’s instructions for 1.5 hours in the dark. Cells were then fixed with 3.2% PFA for 20 minutes in the dark. Cells were washed and resuspended in FACS buffer using a cell scraper and analyzed on a LSRII analyzer as previously described (29). Cells were gated on size, singlets, mCherry (infected)+ and green (dye loaded)+, then analyzed for % of population positive for blue signal.

### Humanized Mouse Infections

NSG mice (NOD.Cg-Prkdc^scid^ Il2^rgtm^1Wjl/SzJ, 005557) engrafted with CD34+ hematopoietic stem cells derived from umbilical cord blood were purchased from The Jackson Laboratory (Bar Harbor, ME). Successful humanization of each mouse is quantified by the supplier from mouse peripheral blood via flow cytometry using anti- hu-CD45+ and anti-murine CD45+, approximately 2 months post-engraftment. Mice were shipped at 31 weeks post-engraftment, allowing increased tissue engraftment of myeloid cells (unpublished data from Jackson Laboratory). After 7 days of acclimation, mice were injected intraperitoneally with a 1:1 ratio of *S.* Typhi strains; either 2 x 10^5^ CFUs for 2 days, or 2 x 10^4^ CFUs of *S.* Typhi for 5 days. The infected mice were closely monitored for signs of illness, and moribund animals were euthanized. At 2 and 5 d, entire livers and spleens were harvested and homogenized in PBS and plated for CFU on LB+Aromix and LB+Aromix+Kanamycin plates to quantify the ratio of WT to mutant bacteria in each organ. The competitive index (CI) was calculated as a ratio of (mutant/wild-type)output / (mutant/wild-type)input.

### Quantification and Statistical Analysis

Statistical significance of all flow cytometry data, CFU counts, time-lapse replication measurements, LB growth curves, and cell death assays were determined by 2-way ANOVA followed by Tukey’s Honestly-Significant-Difference (Tukey Multiple Comparisons) to calculate multiple pairwise comparisons in Prism v. 8.1.2 (GraphPad). Statistical significance of bacteria counted in cells by confocal microscopy was determined by ordinary one-way ANOVA in Prism v. 8.1.2 (GraphPad). Statistical significance of strain replication defects at 24 h.p.i. was calculated by Wilcoxon signed-rank test compared to a hypothetical mean of 1.0 in Prism v. 8.1.2 (GraphPad). Statistical significance of animal infection data was determined by One-sample T-test to a hypothetical mean of 1.0 in Prism v. 8.1.2 (GraphPad), based on the limited availability of validated, aged, healthy, humanized mice available for experiments.

## Acknowledgements

The authors would like to thank Dr. Manuel Amieva and members of the Monack and Amieva laboratories for valuable discussions. Research reported in this publication was supported by grants R01-AI116059 and R01AI095396 from the National Institute of Allergy and Infectious Diseases, United States (D.M.M.), Paul Allen Stanford Discovery Center on Systems Modeling of Infection (to D.M.M.), Gates Grand Challenge Grant from Bill & Melinda Gates Foundation (to D.M.M.) and the Graduate Research Fellowship Program funded by the National Science Foundation (to M.H.). The content is solely the responsibility of the authors and does not necessarily represent the official views of the National Institutes of Health.

## Supplemental Video Legends

**Supplemental Video 1:** WT strain replication in THP-1 macrophages. Phase = Adherent host cells, Red= *Salmonella*, constitutively expressing mCherry. As bacteria replicate, area of red flouresence expands. Green = *Salmonella* that as recently been phagocytosed at 2 h.p.i., green florescence dilutes out over time as intramacrophage replication occurs (pFCcGi). 20X magnification. Scale: bottom left. Time stamp (h.p.i.): bottom right.

**Supplemental Video 2:** Δ*invA*Δ*ssaV* strain replication in THP-1 macrophages. Phase = Adherent host cells, Red= *Salmonella*, constitutively expressing mCherry. Area of red flouresence remains relatively constant throughout the time course, indicating *Salmonella* are not replicating. Green = *Salmonella* that has recently been phagocytosed at 2 h.p.i, lack of florescence dilution at later timepoints indicates lack of replication (pFCcGi). 20X magnification. Scale: bottom left. Time stamp (h.p.i.): bottom right.

**Supplemental Video 3:** WT strain replication in hMDMs. Phase = Adherent host cells, Red= *Salmonella*, constitutively expressing mCherry. As bacteria replicate, area of red flouresence expands. Green = *Salmonella* that as recently been phagocytosed at 2 h.p.i., green florescence dilutes out over time as intramacrophage replication occurs (pFCcGi). 20X magnification. Scale: bottom left. Time stamp (h.p.i.): bottom right.

**Supplemental Video 4:** Δ*invA*Δ*ssaV* strain replication in hMDMs. Phase = Adherent host cells, Red= *Salmonella*, constitutively expressing mCherry. Area of red flouresence remains relatively constant throughout the time course, indicating *Salmonella* are not replicating. Green = *Salmonella* that has recently been phagocytosed at 2 h.p.i, lack of florescence dilution at later timepoints indicates lack of replication (pFCcGi). 20X magnification. Scale: bottom left. Time stamp (h.p.i.): bottom right.

## Notes

### Competing Interest Statement

The authors have declared no competing interest.

