## Supplemental_Figs for "*Salmonella enterica* serovar Typhi uses two type 3 secretion systems to replicate in human macrophages and to colonize humanized mice"

### Figure 1

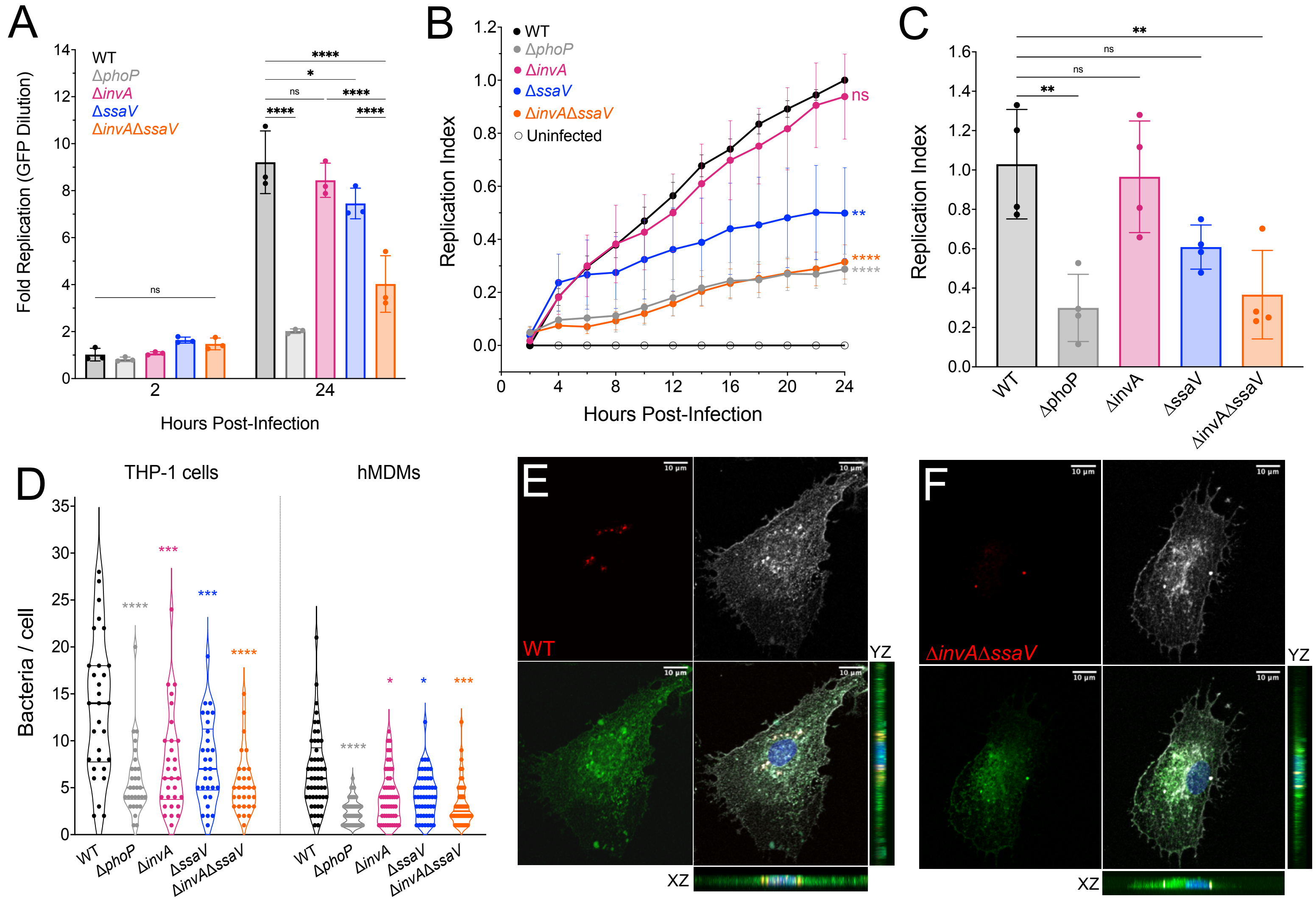
A
Fold Replication (GFP Dilution)
WT
ΔphoP
ΔinvA
ΔssaV
ΔinvAΔssaV
ns
****
*
ns
****
****
****
2
24
Hours Post-Infection
B
Replication Index
WT
ΔphoP
ΔinvA
ΔssaV
ΔinvAΔssaV
Uninfected
ns
**
****
****
Hours Post-Infection
C
Replication Index
**
ns
ns
**
WT
ΔphoP
ΔinvA
ΔssaV
ΔinvAΔssaV
D
THP-1 cells
hMDMs
Bacteria / cell
****
***
***
****
****
*
*
***
WT
ΔphoP
ΔinvA
ΔssaV
ΔinvAΔssaV
WT
ΔphoP
ΔinvA
ΔssaV
ΔinvAΔssaV
E
WT
YZ
XZ
F
ΔinvAΔssaV
YZ
XZ

### Figure 2

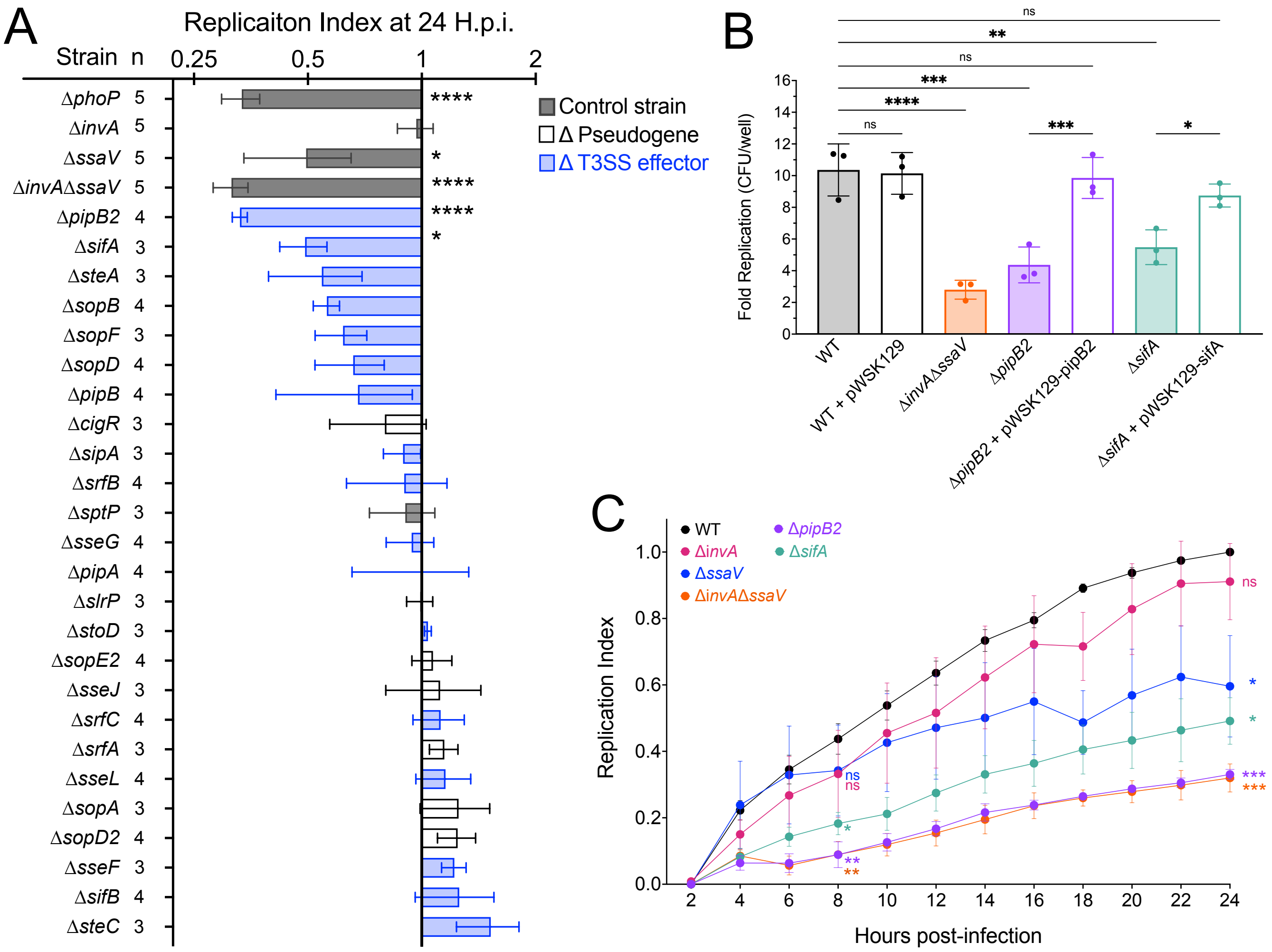

A
Replicaiton Index at 24 H.p.i.
Strain n
0.25
0.5
1
2
ΔphoP 5
ΔinvA 5
ΔssaV 5
ΔinvAΔssaV 5
ΔpipB2 4
ΔsifA 3
ΔsteA 3
ΔsopB 4
ΔsopF 3
ΔsopD 4
ΔpipB 4
ΔcigR 3
ΔsipA 3
ΔsrfB 4
ΔsptP 3
ΔsseG 4
ΔpipA 4
ΔslrP 3
ΔstoD 3
ΔsopE2 4
ΔsseJ 3
ΔsrfC 4
ΔsrfA 3
ΔsseL 4
ΔsopA 3
ΔsopD2 4
ΔsseF 3
ΔsifB 4
ΔsteC 3
Control strain
Δ Pseudogene
Δ T3SS effector
B
Fold Replication (CFU/well)
WT
WT + pWSK129
ΔinvAΔssaV
ΔpipB2
ΔpipB2 + pWSK129-pipB2
ΔsifA
ΔsifA + pWSK129-sifA
ns
C
WT
ΔinvA
ΔssaV
ΔinvAΔssaV
ΔpipB2
ΔsifA
Replication Index
Hours post-infection

### Figure 3

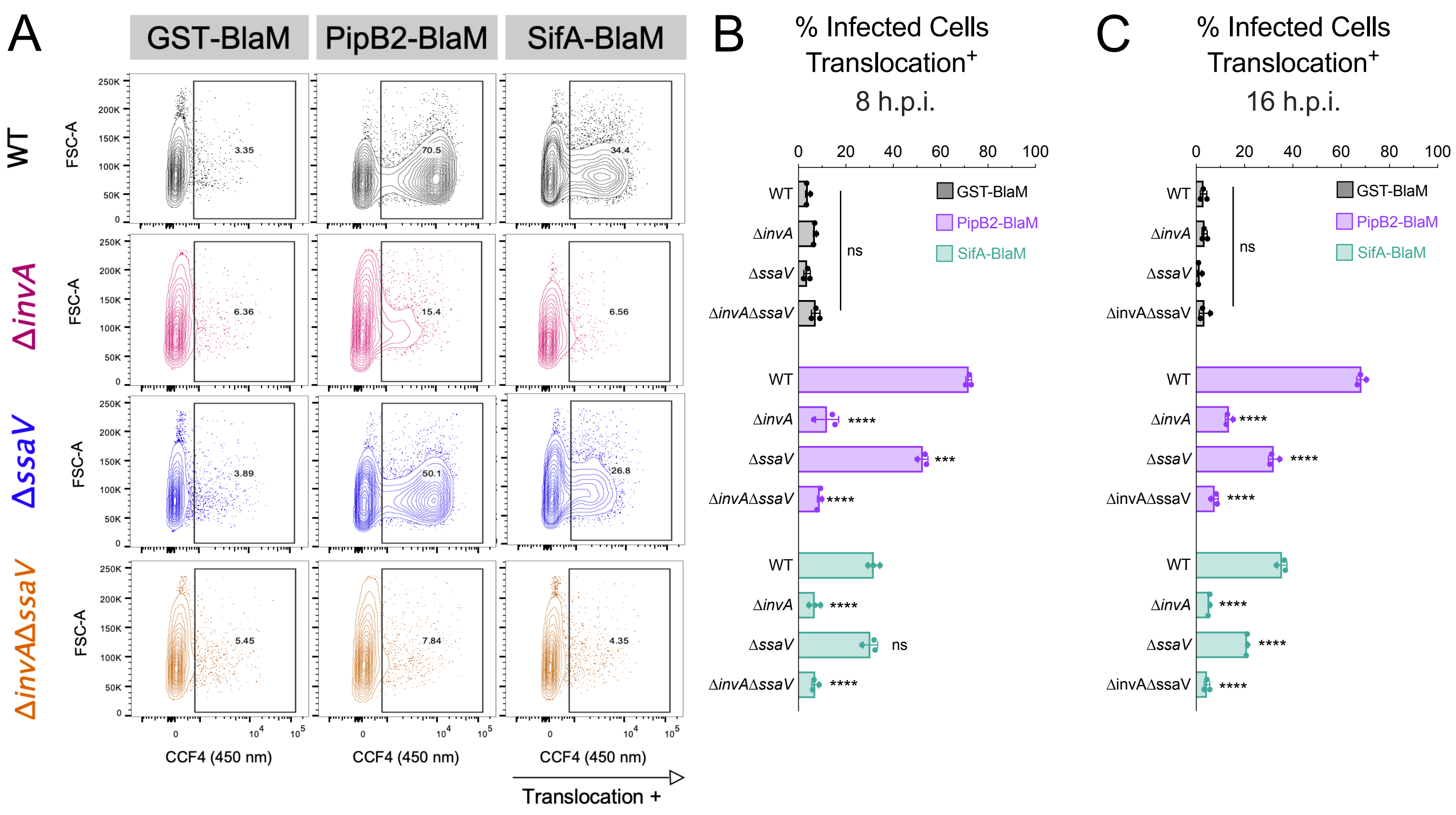

A
GST-BlaM
PipB2-BlaM
SifA-BlaM
WT
ΔinvA
ΔssaV
ΔinvAΔssaV
FSC-A
3.35
70.5
34.4
6.36
15.4
6.56
3.89
50.1
26.8
5.45
7.84
4.35
CCF4 (450 nm)
Translocation +
B
% Infected Cells Translocation+
8 h.p.i.
C
% Infected Cells Translocation+
16 h.p.i.
GST-BlaM
PipB2-BlaM
SifA-BlaM
ns
****
***

### Figure 4

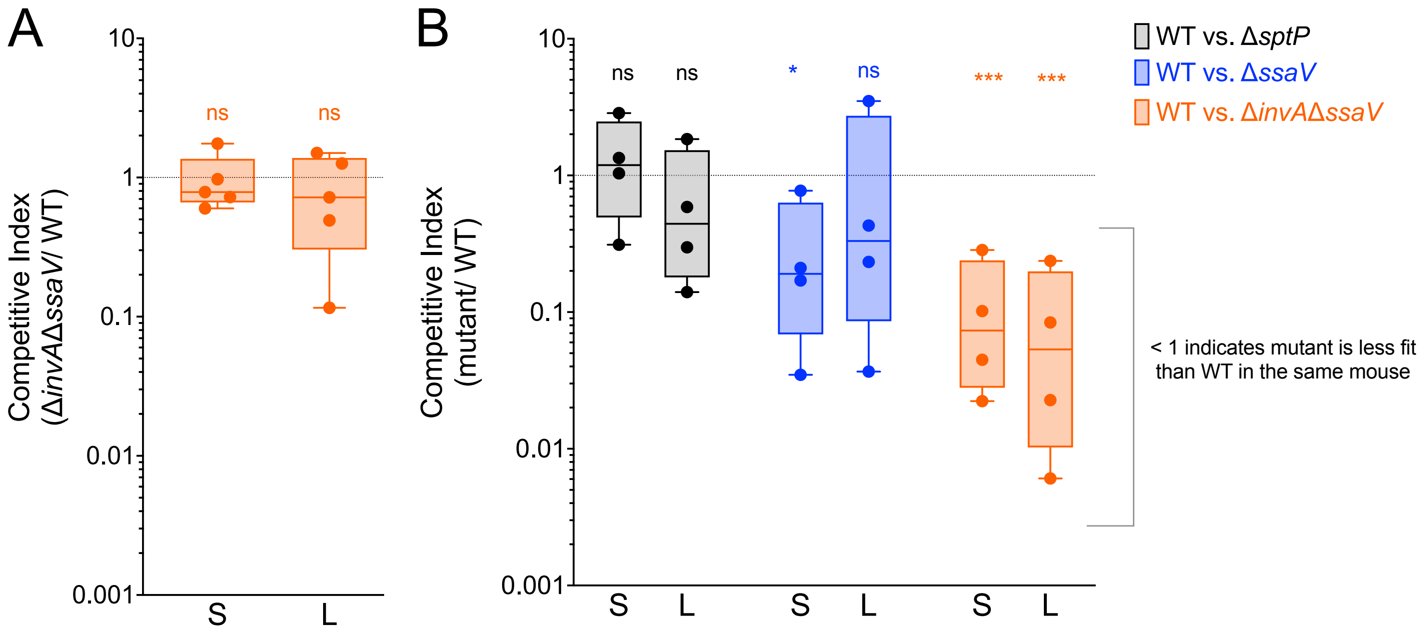
A
B
10
1
0.1
0.01
0.001
Competitive Index
(ΔinvAΔssaV/ WT)
ns
ns
S
L
Competitive Index
(mutant/ WT)
ns
ns
*
ns
***
***
WT vs. ΔsptP
WT vs. ΔssaV
WT vs. ΔinvAΔssaV
< 1 indicates mutant is less fit
than WT in the same mouse

### Supplemental Figure S1

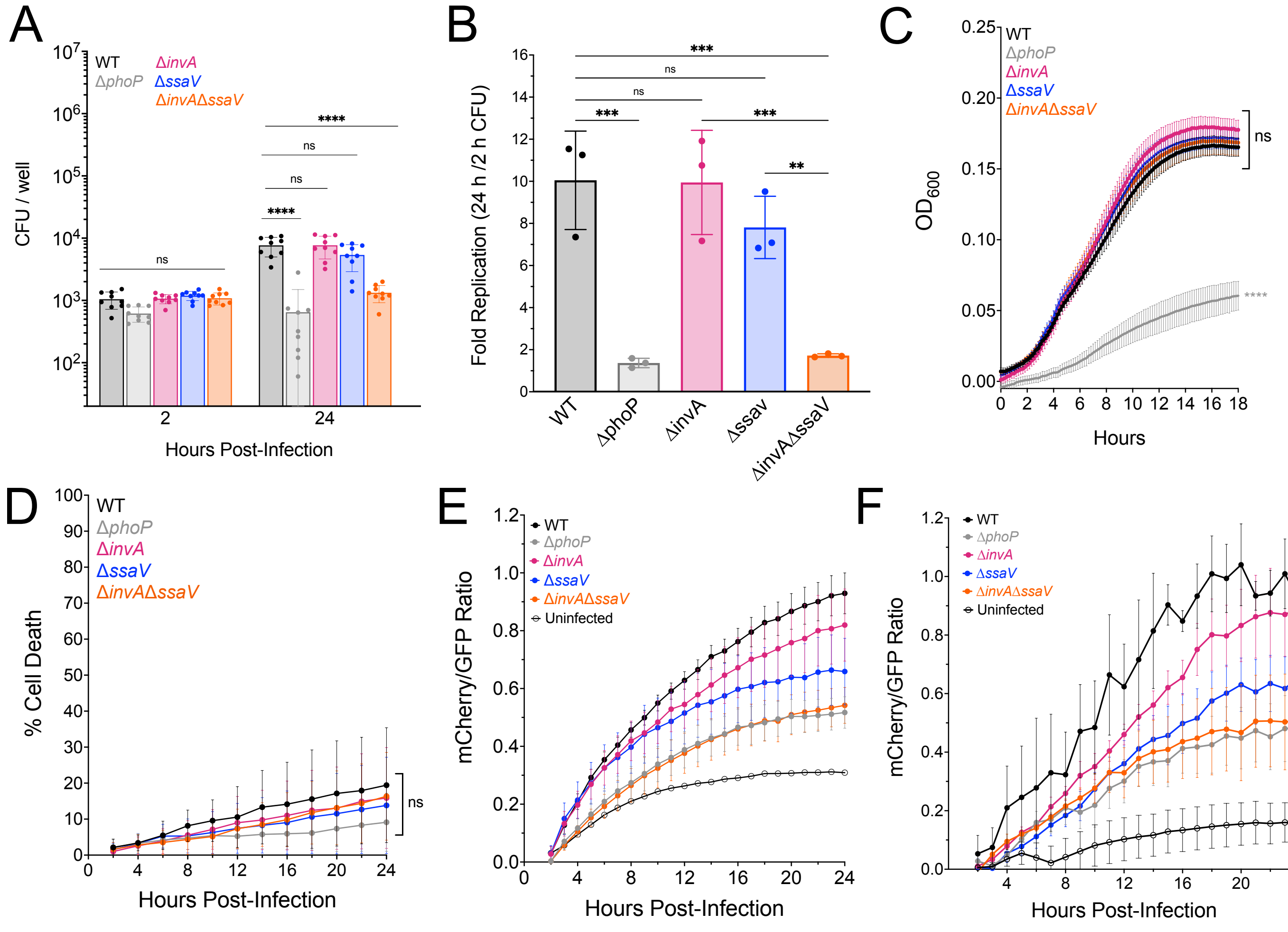


**Figure S1. A.** Colony forming units (CFU) per well of infected THP-1 macrophages. Statistical significance by ANOVA. Bars: mean. Dots: viable bacteria obtained from each well, 3 wells per 1 biological replicate.

**B.** Replication of *S*. Typhi in THP-1 macrophages by CFU/well at 2 and 24 h.p.i. Statistical significance by ANOVA. Dots: biological replicates, each an average of 3 technical replicates. Bars: mean. Error: SD.

**C.** Optical Density (OD) at 600 nm of *S*. Typhi strains in SPI-2-inducing medium (PCN, 0.4 mM $KPO_4$, pH 5.8). Statistical significance by ANOVA. Line: Mean $OD_{600}$of 3 biological replicates every 15 minutes. Error: SEM.

**D.** Percent of $mCherry^+$ (Infected) THP-1 macrophages also positive for Sytox green (Dead) based on automated images taken every hour for 32 hours. At 34 h.p.i. 1% Triton-X added to wells to confirm dye presence and to determined 100% lysis. Statistical significance by ANOVA. Line: Mean of 3 biological replicates. Error: SD.

**E.** Ratio of mCherry/GFP throughout infection of THP-1 macrophages by time-lapse microscopy, prior to subtraction of background signal obtained in uninfected wells. Dots: Mean of 6 biological replicates. Error: SEM.

**F.** Ratio of GFP/mCherry throughout infection of hMDMs by time-lapse microscopy. Data displayed is prior to subtraction of background signal obtained in uninfected wells. Dots: Mean of 4 biological replicates. Error: SEM.

For all; ns = p-value > 0.05, *≤0.05, ** ≤0.01, *** ≤0.001, **** ≤0.0001

### Supplemental Figure S2

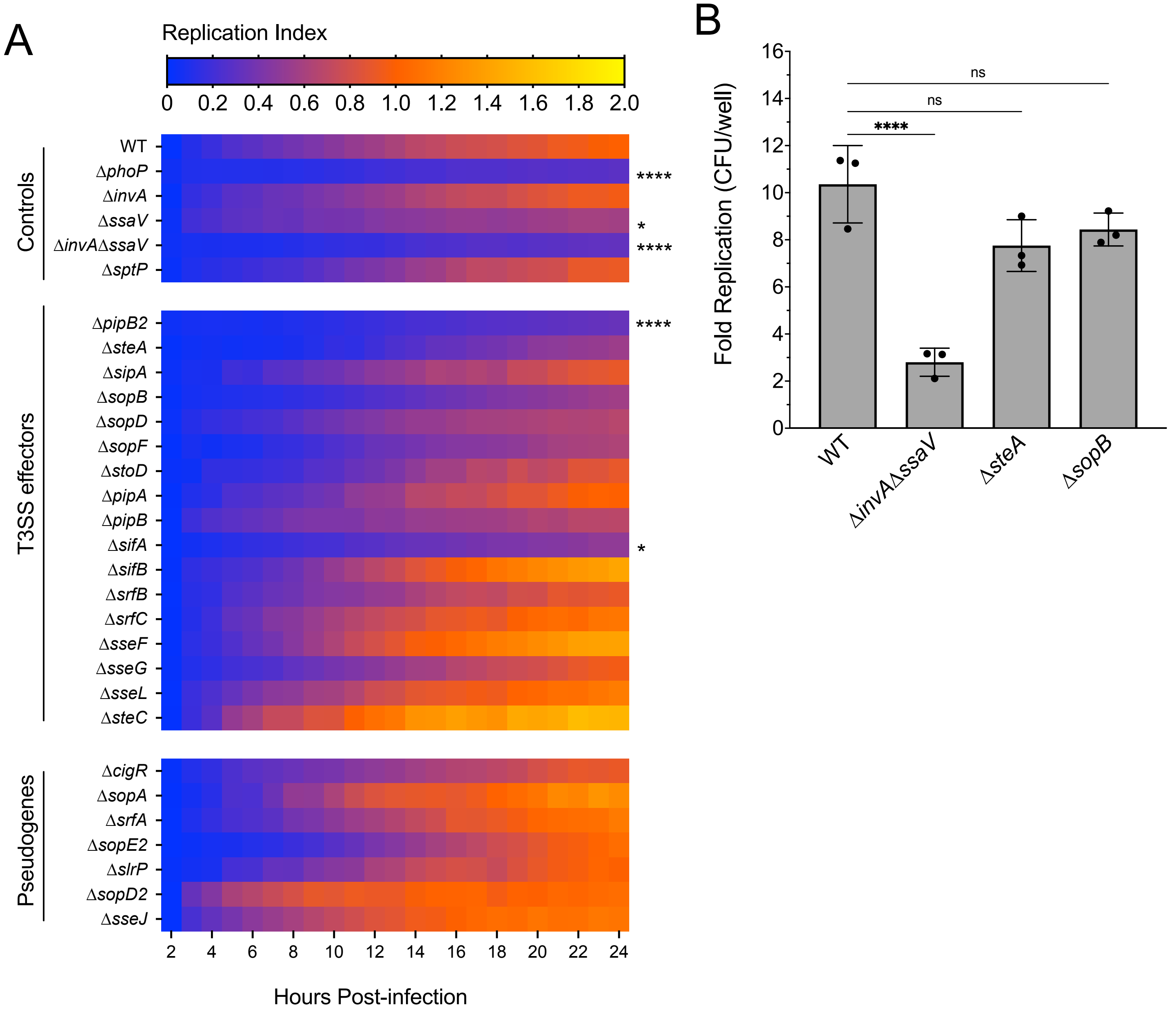


**Figure S2. T3SS-dependent effectors contribute to *S*. Typhi replication in THP-1s.**
**A.** Heat map of replication index over 24 hours for each *S*. Typhi strain in THP-1 macrophages by time-lapse florescence microscopy. Each square: mean of 3-5 biological replicates. Statistical significance at 24 h.p.i. compared to hypothetical mean of 1.0 by Wilcoxon test.
**B.** Replication of *S*. Typhi in THP-1 macrophages by CFU/well between 2 and 24 h.p.i. Statistical significance by ANOVA. Dots: biological replicates, each an average of 3 technical replicates. Bars: mean. Error: SD.

Supplemental Figure S3

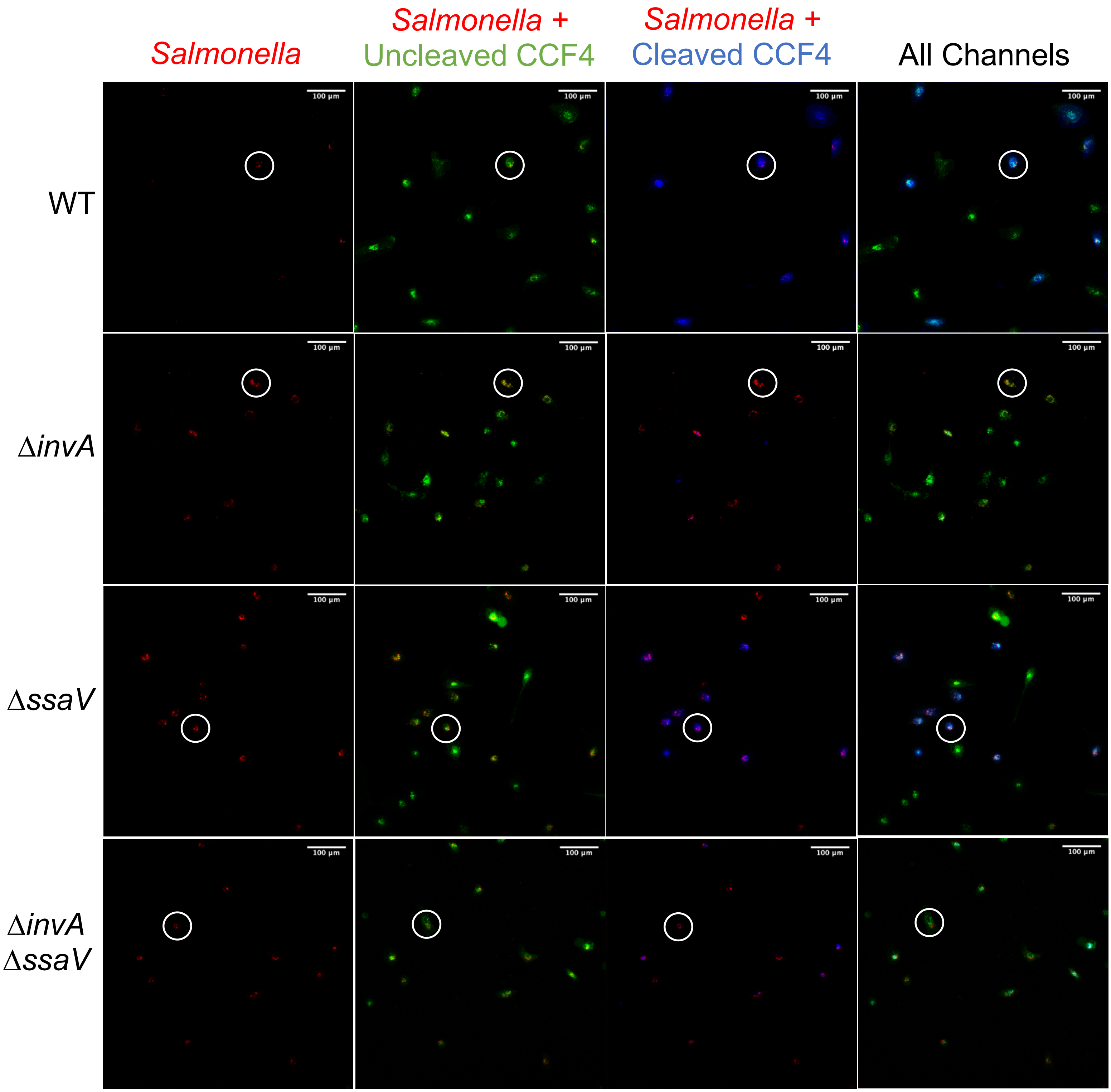


**Figure S3. Representative images of SifA-BlaM translocation in hMDMs 8 hours post-infection**
hMDMs infected with Ty2 strains expressing SifA-BlaM. Infected wells dyed with CCF4, fixed and imaged at 20X, 8 h.p.i. Columns = channels, in order of Red: *Salmonella*, Green: Uncleaved CCF4 dye, indicating hMDM cytosol, Blue: Cleaved CCF4 dye, indicating BlaM in hMDM cytosol, Composite: all three channels overlayed. Rows = *S.* Typhi WT or T3SS knock-out strain. Scale Bar: upper right. White circles indicate a representative example of one infected hMDM in each channel and the composite image of the three channels.

### Supplemental Figure 4

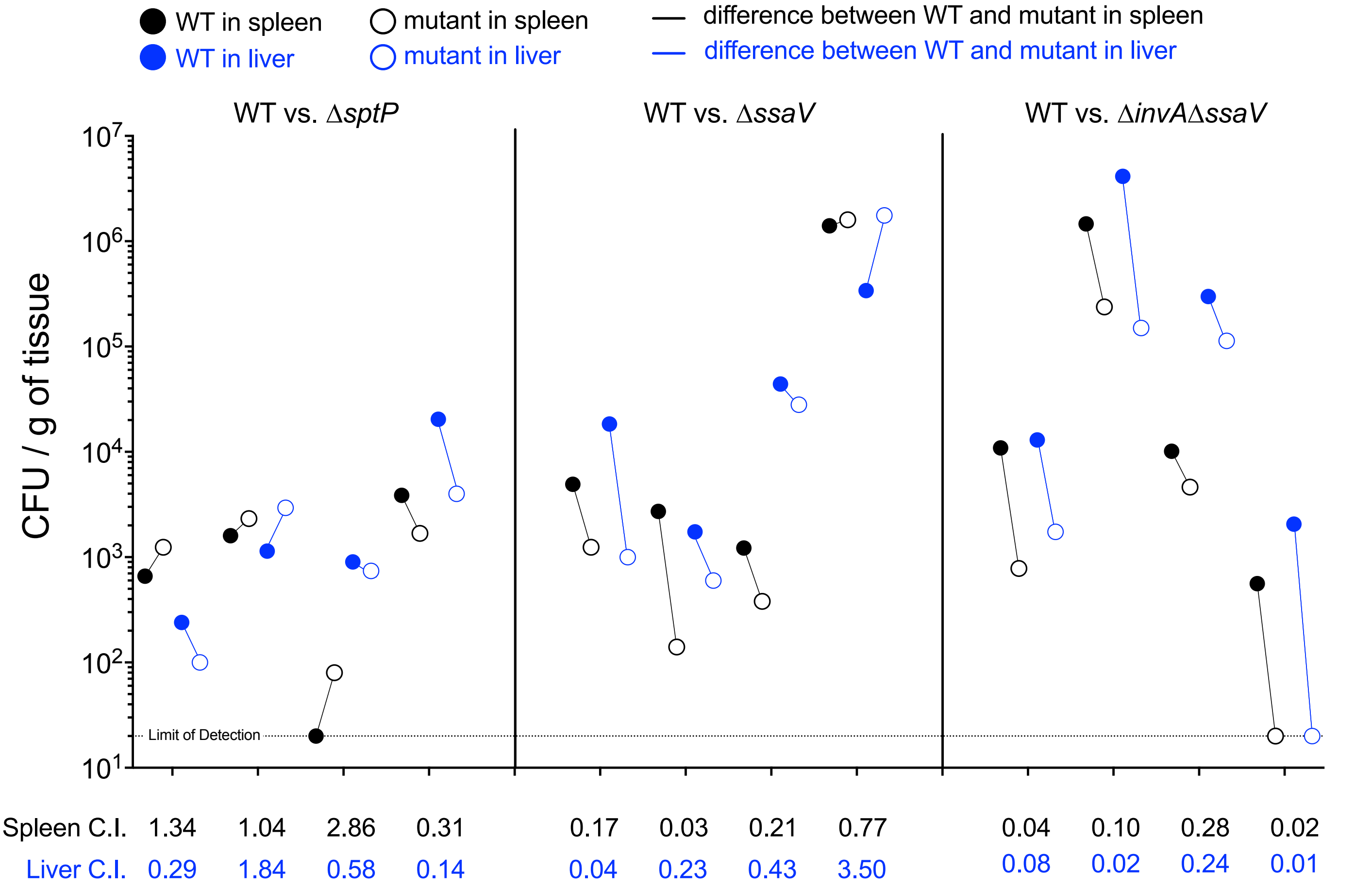


**Figure S4. CFU per gram of tissue in spleen and liver of individual infected mice at 5 days p.i.**
*S*. Typhi burden in whole spleens (black symbols) and livers (blue symbols) sampled for each mouse at 5 days post-infection. Individual mice across X-axis. Y axis indicates CFU per g of homogenized tissue, WT (solid circles) and mutant strain (empty circles) abundance distinguished by kanamycin resistance. Limit of detection of 20 CFU/g indicated by dotted horizontal line. A line connects WT and mutant counts from the same organ.
